# B Cell Tolerance and BCR Signaling Dysregulation in NF155-Mediated Autoimmune Nodopathies

**DOI:** 10.1101/2025.08.10.669569

**Authors:** Bhaskar Roy, Abeer H. Obaid, Zhenjian Wang, Kristof G. Kovacs, Sarah Ohashi, F. Naz Cemre Kalayci, Daniel Joo, Gianvito Masi, Carmina Coppola, Sameeran Das, Amanda L. Hernandez, Lorena Martin-Aguilar, Cinta Lleixà, Richard J. Nowak, Luis Querol, Kevin C. O’Connor

**Author notes:** **Corresponding author:** Bhaskar Roy, Department of Neurology 15 York Street, LCI 916 Yale School of Medicine New Haven, CT, 06510.

## Abstract

**Objective:** Autoimmune nodopathies (AINs) are a group of rare, acquired autoimmune neuropathies with distinct clinical features and the presence of circulating autoantibodies - often of the immunoglobulin G4 (IgG4) subclass - targeting proteins at the node of Ranvier. Defects in B cell tolerance checkpoints have been implicated in several autoimmune diseases. Prior work identified defective B cell tolerance—reflected by a high frequency of self-reactive naïve B cells—in patients with MuSK-positive myasthenia gravis (MG), mediated by IgG4 autoantibodies. Here, we investigated whether tolerance defects exist in neurofascin-155-mediated AIN (NF155-AIN), similar to MuSK+ MG. Additionally, we analyzed B and T cell transcriptomics and interactions at the single-cell level to explore the underlying pathomechanism.

**Methods:** Using a well-established assay, we assessed B cell tolerance fidelity by generating recombinant antibodies from new emigrant (NE) and mature naïve (MN) B cell populations— directly downstream of key tolerance checkpoints—from three NF155-AIN patients, and testing these antibodies for polyreactivity and autoreactivity, thereby determining the frequency of polyreactive and autoreactive B cells. The transcriptome of peripheral blood mononuclear cells (PBMC) was studied, with a special focus on naïve B cells and CD4+ T cells at the single-cell level, along with characterization of cell-cell interactions.

**Results:** NF155-AIN patients have an elevated frequency of polyreactive B cells in the NE (37.4% compared to 9.7% in healthy controls (HCs), p = 0.03) and MN (31.5% compared to 10.5% in HCs, p = 0.03) compartments with increased B cell clones expressing autoreactive antibodies, consistent with a breach in early tolerance checkpoints. We observed abnormal B cell receptor (BCR) signaling characterized by low CD79B, CSK, BLNK, and BTK expression, which may contribute to a breach in B cell tolerance. We also observed evidence of impaired follicular helper T cells (Tfh) and regulatory T cells (Treg), which may limit the normal development and suppression of autoreactive B cells. Moreover, comparative gene expression analysis of B cells and CD4+ T cells from three patients with chronic inflammatory demyelinating polyneuropathy (CIDP) —a related autoimmune neuropathy— confirmed that these differences are largely specific to NF155-AIN, supporting a distinct pathophysiology in this subset.

**Conclusion:** These findings demonstrated a breach in early B cell tolerance checkpoints, defective BCR signaling, and disrupted T cell–B cell interactions in NF155-AIN, all of which may contribute to the development of pathogenic autoreactivity. These immunologic abnormalities appear distinct from those seen in CIDP, supporting NF155-AIN as a unique immunopathologic entity.

## Introduction

Autoimmune nodopathies (AINs) are a rare group of heterogeneous disorders with autoantibodies targeting antigens at the node of Ranvier, including contactin 1 (CNTN1), contactin-associated protein 1 (Caspr1), neurofascin 155 (NF155), neurofascin 186 (NF186).^1–3^ While demyelination in nerve conduction studies is similar between chronic inflammatory demyelinating polyneuropathy (CIDP) and patients with AINs, they have distinct clinical features, and patients with CIDP lack identifiable autoantibodies.^3,4^ Patients with NF155-mediated AIN (NF155-AIN), which is more common in young males, may have a progressive course of symmetric distal-predominant weakness, sensory symptoms, tremors, and ataxia.^1,4^ Additionally, these patients often show poor response to standard immunoglobulin (Ig) therapy but tend to improve with B cell–depleting treatments such as rituximab.^1,3,4^ These NF155 autoantibodies are predominantly of the IgG4 subclass, though other subclasses—including IgG1, IgG2, and IgG3—and antibody isotypes (IgM) can also be detected in some patients. However, clinical features of patients with non-IgG4 predominant NF155 autoantibodies differ from those with predominant IgG4 autoantibodies.^5–8^ Human IgG4 is a distinct antibody subclass with unique structural and functional features. Through Fab-arm exchange, it becomes functionally monovalent, preventing effective cross-linking and immune complex formation. It binds poorly to activating Fcγ receptors and does not efficiently trigger complement activation, limiting its ability to mediate inflammatory responses. As such, IgG4 is generally associated with immune tolerance and chronic antigen exposure rather than immune activation.^9,10^ These characteristics of IgG4 antibodies may contribute to the limited effectiveness of immunoglobulin therapy in NF155-AIN, yet the precise mechanisms driving the disease are still not fully understood. ^9,11^

The generation of a highly diverse B cell repertoire relies on random rearrangement of the variable (V), diversity (D), and joining (J) immunoglobulin gene segments in B cell precursors.^12,13^ While such random arrangement creates antibodies with diverse amino acid sequences that can recognize a broad spectrum of pathogenic antigens, it also produces autoreactive B cells.^13–15^ In healthy individuals, two tolerance checkpoints remove these autoreactive B cells.^14,16^ The first is the central tolerance checkpoint, which occurs in the bone marrow, and is critical in removing self-reactive clones from the early immature to immature B cell stage.^14,16^ The second is the peripheral tolerance checkpoint, which eliminates most remaining autoreactive new emigrant/transitional B cell clones prior to becoming mature naive B cells.^14,16,17^

Dysfunctional B cell tolerance checkpoints, which allow autoreactive B cells to persist, have been implicated in many autoimmune diseases, but the integrity of the B cell tolerance checkpoints has not been tested in NF155-AIN. Among diseases involving defects in B cell tolerance checkpoints, myasthenia gravis (MG) with autoantibodies against muscle-specific tyrosine kinase (MuSK+ MG) - another IgG4-mediated condition - is of particular interest.^16,18–20^ This is because both NF155-AIN and MuSK+ MG exhibit distinct clinical phenotypes compared to the more prevalent disease subtypes—(CIDP for NF155-AIN; MG with autoantibodies against acetylcholine receptor (AChR+ MG) for MuSK+ MG)—and both respond better to B cell depletion with rituximab than AChR+ MG or CIDP.^21–23^ Both MuSK+ and AChR+ MG subtypes exhibit defects in B cell tolerance; however, our previous work showed that MuSK+ MG patients uniquely demonstrated a higher frequency of polyreactive new emigrant B cells in MuSK+ MG patients compared to those with AChR+ MG.^16,18^ Furthermore, anti-MuSK monoclonal antibodies isolated from MuSK+ MG patients retained strong MuSK binding even after mutations acquired through affinity maturation were reverted to their original germline sequences, which emulate the naïve B cell receptor. This suggests that defects in tolerance checkpoints allowed early MuSK-specific B cell precursors to escape deletion, interact with MuSK, undergo somatic hypermutation, and contribute to disease pathogenesis.^24^ Given these findings in MuSK+ MG, we sought to determine whether a similar breach of tolerance checkpoints occurs in NF155-AIN and whether the underlying pattern resembles that of MuSK+ MG. Furthermore, we analyzed the transcriptomic profiles of naïve B cells and CD4+ T cells at the single-cell level and examined their ligand-receptor interactions to gain insight into the underlying mechanisms of tolerance checkpoint defects in NF155-AIN. We also compared our findings of NF155-AIN with three patients with CIDP, and three patients with MuSK+ MG to further explore the specificity of our observations.

## Methods

### Study approval, subjects, and specimens

Deidentified peripheral blood mononuclear cells (PBMCs) from patients with NF155-AIN, isolated by CPT tubes or standard Ficoll-gradient protocol were collected at the Hospital de la Santa Crue I Sant Pau in Spain (Dr. Luis Querol) and Yale School of Medicine. The presence of NF155 was confirmed by cell-based assay (CBA) or ELISA from a CLIA (Clinical Laboratory Improvement Amendments)-certified laboratory. Details of previous immunosuppressive therapies were reviewed for all NF155-AIN patients to ensure that none received any immunosuppression, which can affect B cell generation and maturation before sample collection, particularly B cell depletion therapy. PBMC samples of age and gender-matched healthy controls (HCs) and patients with CIDP and MuSK+ MG were obtained from our biorepository.^18,25^ All subjects recruited in the B cell tolerance checkpoint assay were tested for PTPN22*R620W, a polymorphism associated with dysfunctional central and peripheral tolerance regulation, and all were confirmed to be negative. Sample collection and all experiments were approved by the Institutional Review Board at the Yale School of Medicine.

### Cell staining and sorting

Single B cells from PBMCs were isolated as previously described.^18,25^ Briefly, CD20+ B cells were isolated with magnetic beads (Miltenyi). Additional fluorescence-activated cell sorting on a FACSAria flow cytometry (BD) was then performed to sort single B cells, namely new emigrant (NE)/transitional (CD19+, CD21^low^, CD10+, IgM^high^, CD27-) and mature naïve (MN) B cells (CD19+, CD21+, CD10-, IgM+, CD27-), into 96-well PCR plates. Sorted plates were immediately frozen on dry ice and stored at −80^◦^C prior to complementary DNA synthesis.

### Recombinant antibody production

Central and peripheral B cell tolerance defects were assessed using an established protocol that measures the frequency of polyreactive B cells downstream of the two checkpoints.^18,25^ Higher frequencies of polyreactive B cells indicate a compromised ability to remove autoreactive B cell clones. Specifically, NE/transitional B cells and MN B cells were isolated from peripheral blood and sorted into individual wells. The complementary DNA (cDNA) synthesis from the single sorted B cells, heavy and light chain PCR amplification, cloning strategy, expression vectors, and antibody expression and purification were conducted as previously described.^18,25^ The IgBLAST tool^26^ on the NCBI website or the IMGT/V-QUEST tool (available on the International ImMunoGeneTics (IMGT) information system website^27^) were used to assess the germline fidelity of the immunoglobulin sequences amplified from single B cell clones. Each recombinant antibody was cloned and expressed in duplicates.

### ELISA and indirect immunofluorescence staining

Recombinant IgG in cell culture supernatants were tested for polyreactivity on 96-well ELISA microplates coated with double-stranded DNA (dsDNA) (Sigma Aldrich), lipopolysaccharide (LPS) (Sigma Aldrich), or recombinant human insulin (Sigma Aldrich) as previously described.^18,25^ Antibodies were considered polyreactive if they recognized all three antigens (dsDNA, insulin, and LPS) with cut-off of ∼0.5 at OD405nm for positivity. Highly polyreactive ED38 was used as a positive control.^18,25^ For indirect immunofluorescence assays, HEp-2 cell-coated slides (Bion Enterprises) were incubated in a moist chamber at room temperature with purified recombinant antibodies at 20 μg/ml and detected with FITC-conjugated goat anti-human IgG.

### Single-cell transcriptomics analysis

Cells were prepared for single-cell analysis following the protocol developed by 10x Genomics.^28^ Isolated PBMCs were enriched for CD45-positive cells using CD45 MicroBeads (Miltenyi Biotec). AIN sample 1, HC sample 1, and MuSK+ MG samples 1 and 2, were enriched for B cells using immunomagnetic negative isolation (STEMCELL technologies). Dead/apoptotic cells were removed using Annexin V immunomagnetic depletion (STEMCELL technologies), and cells were filtered to remove debris and cell clusters. Chromium 10x single cell 5 prime library V2 or V3 was used and sequenced through Illumina NovaSeq for 150 × 150 paired-end reads. Low-quality and dead cells (gene count <200 or >3500 or percent mitochondrial gene >5%), and doublets were excluded from the analysis, and each sample was individually assessed for quality control.^29^ Further filtering, normalization, scaling, sample integration, and principal component analysis were performed using the standard protocol from the Seurat platform (version 5).^30^ For identifying the appropriate resolution for clustering, we used the Clustree package.^31^ For integration, we used canonical correlation analysis (CCA) from the Seurat platform. We used the Azimuth platform for cell annotation, which was also compared to the SingleR platform to confirm accurate annotation.^32,33^ We identified T regulatory (Treg) cells and T follicular helper (Tfh) cells manually using appropriate gene expression signatures (Treg: FOXP3, IL2RA, CTLA4, IKZF2, Tfh: CXCR5, PDCD1, BCL6, ICOS, MAF, IL21) in addition to automated annotation. When applicable, subsets of individual cell subtypes (either B cells or CD4+ T cells) were analyzed after re-clustering.

For differential gene expression, we also used the pseudobulk analysis (gene expression profile per sample and cell type) to reduce false positive associations using DESeq2, which estimates variance across samples, when applicable.^34,35^ We performed pathway enrichment analysis to understand the biological functions over-represented by these genes using both overrepresentation analysis and gene set enrichment (GSE) analysis.^36^ We used the web-based G Profiler platform and the cluster profiler package for pathway enrichment analysis in R.^37,38^ We mainly focused on the Gene Ontology (GO) and Reactome pathways.

### Cell-cell communication analysis

To systematically infer cell-cell communication, scRNA-seq data were integrated with prior ligand-receptor interaction database CellChatDB using the CellChat platform.^39^ CellChat identifies differentially over-expressed ligands and receptors for each cell group, quantifies the communication probability between two interacting cell groups (based on the average expression values of a ligand by one cell group, and that of a receptor by another cell group and their cofactors). Then, CellChat infers significant interactions using permutation tests, and an intercellular communication network of a signaling pathway is computed by summarizing the communication probabilities of all associated ligand-receptor pairs. Network centrality analysis summarized each cell population-associated outgoing and incoming communication probabilities across all signaling pathways. To examine the difference between cell-cell communication between AIN and CIDP, they were compared against healthy controls.

### Statistics

Mann-Whitney and Kruskal-Wallis tests were used to compare between group/groups as applicable. False discovery rate analysis incorporated in relevant Seurat or CellChat packages was utilized for multiple comparisons.

## Results

### Study design and approach

We employed a well-described approach to assess the fidelity of the tolerance checkpoints guiding B cell development.^18,25^ We measured the frequencies of polyreactive and autoreactive BCRs expressed in the NE and MN B cells, downstream of the central and peripheral B cell tolerance checkpoints, respectively, which allowed us to assess the extent of negative selection through these early selection steps. We sorted the NE and MN B cells into single wells, and generated recombinant monoclonal antibodies (rAbs) from each single cell, as a means to represent their BCR, which were then tested for reactivity against a panel of defined antigens, expression of anti-nuclear antibodies (ANA), and compared against healthy controls with properly established tolerance.^17^ These assays have been performed by independent groups and have identified B cell tolerance checkpoint defects in patients with primary immunodeficiency disorders and other autoimmune diseases.^16,40,41^ The approach produces a large effect size when tolerance is compromised; therefore, we recruited three patients with NF155-mediated AIN (53.3 ± 9.3 years, two males), and compared them against four healthy controls (39.5 ± 12.4 years, two males, three of the healthy control data were used in our previous publications). There was no significant difference in terms of age between the groups (p-value 0.23).^18,25^ All the patients with NF155-AIN had distinct clinical presentation, with evidence of demyelination in electrodiagnostic studies, and none of them were treated with B cell depletion prior to sample collection (**Table S1**). A total of 81 unique rAbs from the NE (40) and MN (41) compartments of the three NF155-AIN patients were cloned and expressed in duplicates and compared against 70 NE and 64 MN BCRs from the healthy controls (**Table S2**).

### Early tolerance checkpoint fidelity is compromised in NF155-AIN patients

We examined the frequency of B cell clones expressing polyreactive antibodies. Among the recombinant antibodies (rAbs) from the NE B cells from NF155-AIN, 37.4 ± 11.5% were polyreactive and bound to all three antigens (dsDNA, insulin, and LPS), compared to only 9.7 ± 4.2% of polyreactive NE B cell clones in HCs (p-value 0.03) (**Fig. 1A, B)**. Similarly, 31.5 ± 2.5% of rAbs from the mature naïve B cells from NF155-AIN patients were polyreactive compared to 10.5 ± 3.3% in HCs (p-value 0.03) **(Fig. 1C, D)**.

**Figure 1.**
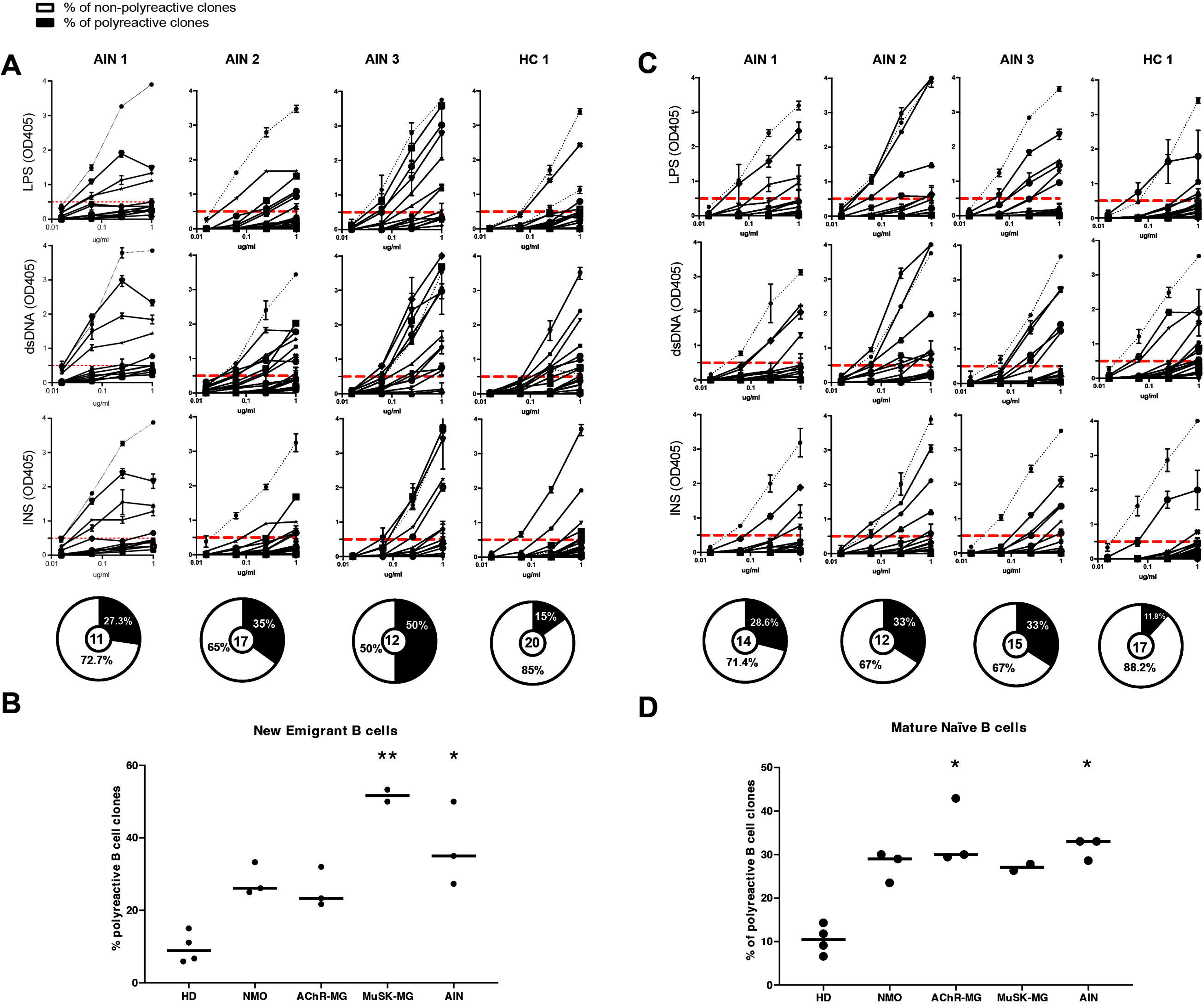
Early B cell tolerance is compromised in patients with NF155-mediated AIN. Recombinant antibodies (rAbs) derived from A) new emigrant (NE)/transitional B cells, and C) mature naïve (MN) B cells of patients with NF155-AIN are polyreactive when tested by ELISA for reactivity against LPS, dsDNA, and insulin (one representative healthy control (HC) data shown). The dotted black line represents the highly polyreactive mAb, ED38, that was used as a positive control, while the dotted horizontal red line marks the positive reactivity cut-off at OD405 reading of 0.5. Pie charts represent the total number of recombinant antibodies tested and the percentage of which displayed polyreactivity (colored in black). B, D) Polyreactive antibody frequencies in AIN, healthy controls, myasthenia gravis, and neuromyelitis optica cohorts in B) new emigrant, and D) mature naïve compartments. The frequency of polyreactive antibodies was plotted for each subject along with the mean for each subject group. Significant statistical differences are shown. [AIN: NF155 mediated AIN, NMO: Neuromyelitis optica, AChR-MG: Myasthenia gravis with autoantibody against acetylcholine receptor, MuSK-MG: Myasthenia gravis with autoantibody against muscle-specific tyrosine kinase, LPS: lipopolysaccharide, dsDNA: double stranded DNA, OD405: optical density at 405 nm]

We examined the frequency of B cell clones expressing antinuclear antibodies (ANA) using HEp-2 cell–coated slides. Overall, 26 ± 3.6% of recombinant antibodies (rAbs) from NE B cell clones in NF155-AIN patients showed ANA reactivity, compared to 9.1% in healthy controls (HCs). Similarly, 24.8 ± 4.5% of rAbs from MN B cell clones in NF155-AIN patients were ANA-positive, while none were detected in HCs. Various ANA patterns, including nuclear, cytoplasmic, and nuclear plus cytoplasmic, were observed (**Fig. 2**). These collective findings confirm defects in early B cell tolerance checkpoints in NF155-AIN.

**Figure 2.**
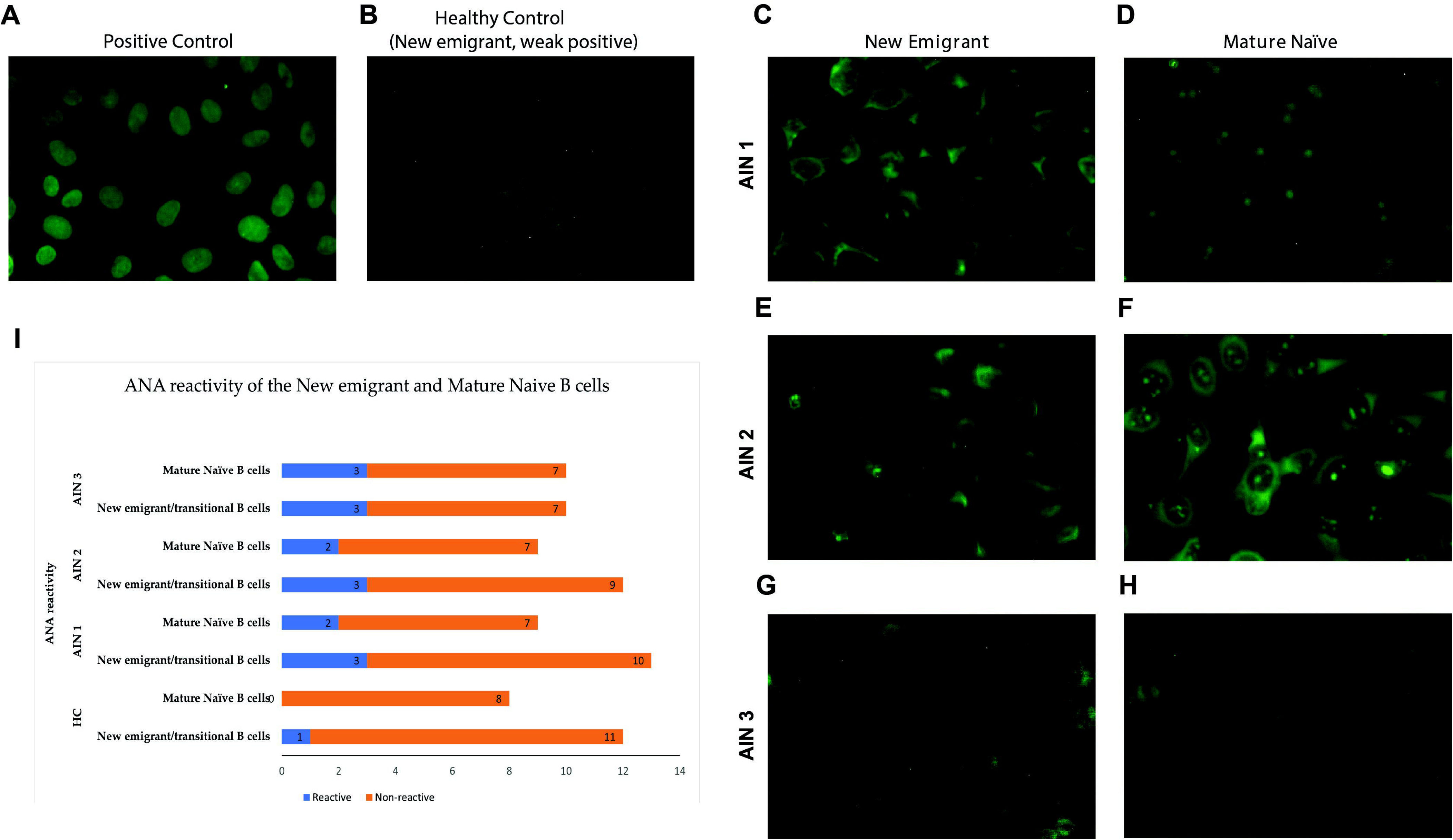
Anti-nuclear antibody (ANA) expressing B cell clones (based on reactivity of recombinant antibodies (rAbs) from new emigrant and mature naïve B cells from patients with AIN and one representative healthy control. Figure shows representative images of A) positive control (positive serum provided by the manufacturer), B) a weak positive new emigrant B cell clone from the healthy control, C-E-G) positive new emigrant B cell clones from AIN patients, and D-F-H) positive mature naïve B cell clones from AIN patients. Nuclear, cytoplasmic, nuclear plus cytoplasmic patterns were prominently noted. I) Bar diagram shows the number of ANA expressing clones (blue) and non-ANA expressing clones (orange) in AIN patients and a healthy control. [ANA: anti-nuclear antibody, AIN: neurofascin 155 mediated autoimmune nodopathy, HC: healthy control]

### Transcriptomics analysis at the single-cell level

B cell tolerance checkpoints are complex processes that occur at multiple stages of B cell development. While central tolerance in the bone marrow is primarily B cell–intrinsic, regulatory T cells (Tregs) are essential for enforcing peripheral tolerance and preventing autoimmunity. In the periphery, CD4+ T cells—particularly follicular helper T cells (Tfh)—play a critical role in B cell activation, differentiation, and antibody production through antigen-specific B–T cell interactions. Therefore, we sought to examine both the B and CD4+ T cell compartments to characterize their transcriptomic profiles at the single-cell level, as a means to gain insight into the potential contributors to tolerance checkpoint breaches and the underlying pathophysiology of NF155-AIN.

For RNA transcriptomics analysis at the single-cell level, PBMC samples from four patients with NF155-AIN were used (age 49 ± 15.7 years, three males). We used seven HCs (48.9 ± 14.6 years, five males), three CIDP patients (54.7 ± 13.1 years, one male), and three MuSK+ MG patients (age 55.7 ± 9.3 years, three women) as controls. There were no significant differences between the groups regarding age (NF155-AIN vs. HC, p-value 0.93; NF155-AIN vs. CIDP, p-value 0.82). All patients with NF155-AIN had typical presentations, and all CIDP patients fit the electrodiagnostic criteria of possible CIDP based on the EAN/PNS diagnostic criteria.^42^ All MuSK+ MG patients had confirmed autoantibodies, and two of them were treated with B cell depletion therapy over a year ago. Details of clinical data and previous therapies are presented in **Supplementary Table 1**.

Since samples were run at different time points, we ensured the proper integration of samples, followed by cell annotation, to address batch effects (**Fig. 3A, Fig. S1A-B**). There were biological variations in terms of the percentage of cell types between NF155-AIN and HC, but overall, they were comparable (**Fig. 3B**).^35,43^ Furthermore, we subset cells of interest (B cells and CD4+ T cells) and reclustered them to perform a comprehensive analysis of transcriptional states (**Fig. 3C** for B cells, **Fig. S1C-D**). We also examined the conserved markers in different subtypes of B cells between conditions (**Fig. 3D**) and examined the top five feature genes in different clusters (**Fig. S2**). As expected, a higher percentage of plasmablasts expressed IgG4 in patients with NF155-AIN, reinforcing the accuracy of cell clustering and annotation. We observed overall reduced expression of CD22, FCER2 (CD23), CD24, and SELL (L-selectin) across different subtypes of B cells, which are important for normal B cell development (**Fig. 3D**).

**Figure 3.**
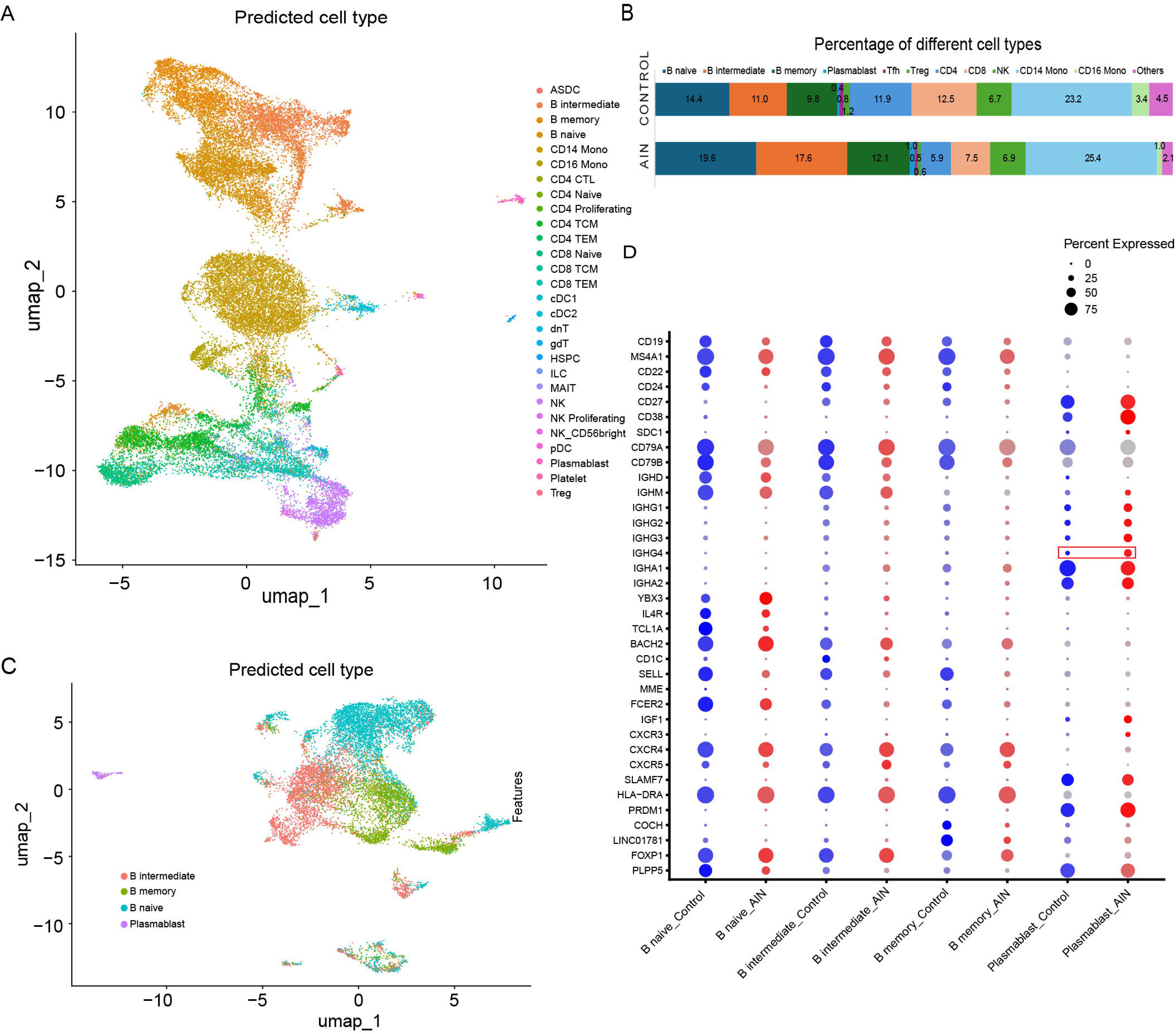
A) Uniform manifold approximation and projection (UMAP) of peripheral blood mononuclear cells (PBMCs) in AIN (n=4) and healthy controls (n=7) showing clustering of different cell populations (each population is indicated by a distinct color). B) Percentage of different cell populations among AIN patients and healthy controls. C) UMAP representation of clusters of different B cell subtypes. D) Expression of selected conserved B cell markers between AIN patients and healthy controls. The red rectangle outlines the percentage of B cells expressing immunoglobulin (Ig) G subclass 4 (IgG4) in AIN patients and healthy controls. [AIN: neurofascin 155 mediated autoimmune nodopathy, HC: healthy control]

### A distinct signature of naïve B cells in NF155-AIN patients

We examined the differential gene expression in B naïve cells (741 upregulated genes and 3118 downregulated genes in NF155-AIN), and some key highly significant genes (p-value < e^-50^) with Log2 fold change of 1 or higher are represented in a volcano plot in **Fig. 4A**. GSE (**Fig. 4E**) and overrepresentation analysis (**Fig. 4F, G**) showed that pathways associated with B cell activation, differentiation, B cell homeostasis, T helper cell differentiation, and intracellular transport were significantly different between NF155-AIN and control. Among the differentially expressed genes, CD79B, a critical component of the BCR complex, was reduced in patients with NF155-AIN (**Fig. 4A, C**). While LYN and NFKB1 were over-expressed, several downstream genes related to BCR signaling, including SYK, CSK, BTK, and BLNK, were downregulated, suggesting abnormal BCR signaling in NF155-AIN. Some genes related to intracellular transport, including SMAP2 and SNX9, were upregulated in patients with NF155-AIN. Wiskott-Aldrich syndrome (WAS) gene, expressing a multidomain protein involved in signal transduction to the actin cytoskeleton, was also downregulated in patients with NF155-AIN, and WAS deficiency is strongly associated with autoimmunity (**Fig. 4A-D, Figure SD**).^44^ Expression of TCL1A, an important factor for B cell regulation and differentiation, was lower in naïve B cells in NF155-AIN patients (**Fig. 4A**).

**Figure 4.**
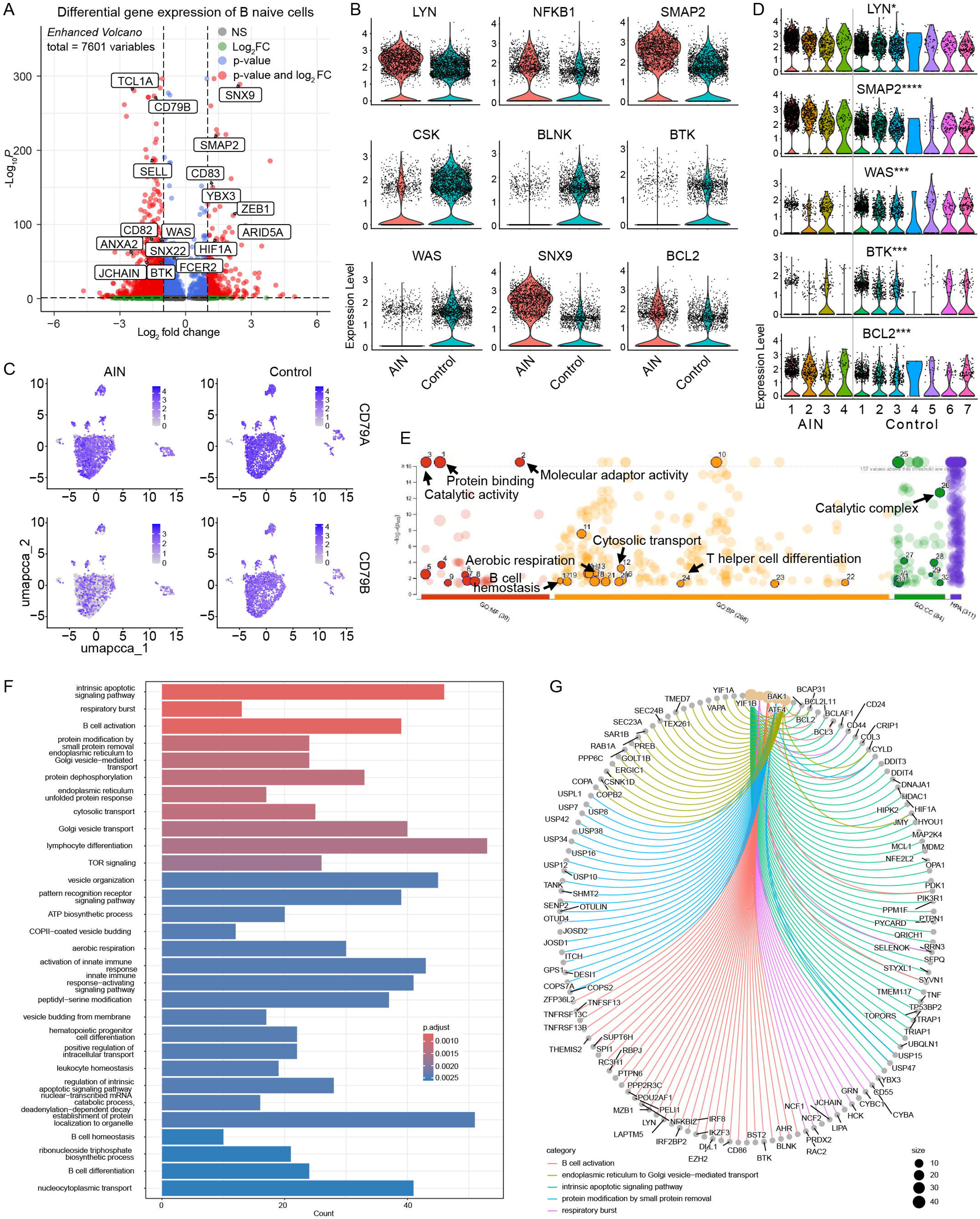
A) Volcano plot shows differential gene expression between B naïve cells from AIN patients and healthy controls. The dotted horizontal line shows the p-value cut off and vertical lines show log2-fold change in gene expression. B-D) Expression of some genes between groups and individual sample levels. E) Manhattan plot of gene set enrichment shows pathways related to molecular function (MF), biological process (BP), and cellular components (CC) based on gene ontology (GO) database, and Reactome pathway. p-values were capped at e^-16^ for graphical representation and shown as a dotted line. Pathways of interest are annotated. Images were created using g:Profiler platform. F) Bar plot shows top relevant pathways with gene count differentiating AIN patients and healthy controls. G) Network plot shows different genes which were expressed differentially and the associated pathways. [AIN: neurofascin 155 mediated autoimmune nodopathy, HC: healthy control, **** p < 0.001, *** p < 0.005, ** p < 0.01, * p < 0.05, for panel B, all changes were significant at p<0.001]

While this work focused on B naïve cells, we also examined B intermediate cells (transitional cells between naïve and memory B cells), and memory B cells; similar abnormalities associated with B cell activation and differentiation were noted in both compartments (**Fig. S4, S5**). Apart from low CD79B, SELL, and upregulated SNX9 expression, we also noted low expression of NCF1, FCMR, FCRLA, and PTPN6 in B intermediate cells. NADPH oxidase complex (NOX2) genes, such as NCF1, which can modulate endosomal Toll-like receptor (TLR signals), and have been implicated in systemic lupus erythematosus and in breaks of B cell tolerance.^45^ Similarly, low expression of FCMR, an Fc receptor specific for pentameric IgM, can lead to impaired B cell homeostasis and increased autoantibody production.^46^ FCRLA, which is engaged in antibody assembly but is not indispensable for antigen-specific humoral immune responses, was also reduced in NF155-AIN.^47^ PTPN6 is also critical for B cell development and hemostasis.^48^ In the memory B cell compartment (**Fig. S5**), SELL, PTPN6, Lat 2, POLD4, and ANXA2 were downregulated. While antibodies to ANXA2 have been detected in Lyme disease and antiphospholipid syndrome (APS), the exact role of decreased expression of ANXA2 in autoimmunity is unclear.^49,50^ Lat2 may promote the survival of autoreactive B cells.^51,52^ POLD4 is a subunit of DNA polymerase δ, and could impair error-free repair during somatic hypermutation and class switch recombination, and such an error in base excision repair has been implicated in autoimmunity.^53^ SCIMP, a transmembrane adaptor protein involved in MHC II signaling, was also downregulated in patients with NF155-AIN.^54^ Overall, these findings suggest an abnormal B cell homeostasis promoting autoimmunity and aberrant autoantibody production in NF155-AIN.

### Impaired CD4+ T cell function in NF155-AIN patients

Given the critical role of CD4+ T cells in B cell maturation, we examined the differences in the CD4+ T cell population from NF155-AIN and HC (**Fig. 5**). Pathways involved in CD4+ T cell differentiation, activation, B cell activation, and cellular respiration were impaired in NF155-AIN (**Fig. 5A-B, F**). We noted low IL-32 and Foxp3 expression in T regulatory cells in patients with NF155-AIN, which is of particular significance, as low expression of the proinflammatory cytokine IL-32 leads to reduced Foxp3 expression, which is a lineage specification factor for Treg cells (**Fig. 5C-E**).^55^ High Foxp3 expression is essential for phenotypic stability and regulatory function of Tregs.^56^ Bach2 is also a critical transcriptional factor in the formation and function of CD4+ T cell lineages and Treg cells; and it was upregulated in NF155-AIN.^57,58^ We also noted a higher percentage of Tfh cells expressing MAF, a transcription factor, concomitantly with BCL6, which are critical for Tfh differentiation,^59,60^ in healthy controls compared to patients with NF155-AIN. Interferon regulatory factor 1 (IRF1) was upregulated in NF155-AIN, and higher rates of single nucleotide polymorphisms (SNPs) in IRF1 have been reported in multiple sclerosis (MS) and in IRF4 for rheumatoid arthritis.^61^ We also noted differential expression of CD55, TSPO, and RBFOX2, which have been implicated in immune dysfunction, but the exact impact of their altered expression in CD4+ T cells is yet to be determined. Overall, these findings suggest that, in addition to the B cell tolerance checkpoint defect in NF155-AIN, there is impaired CD4+ T and B cell interaction, which can further lead to abnormal B cell development.

**Figure 5.**
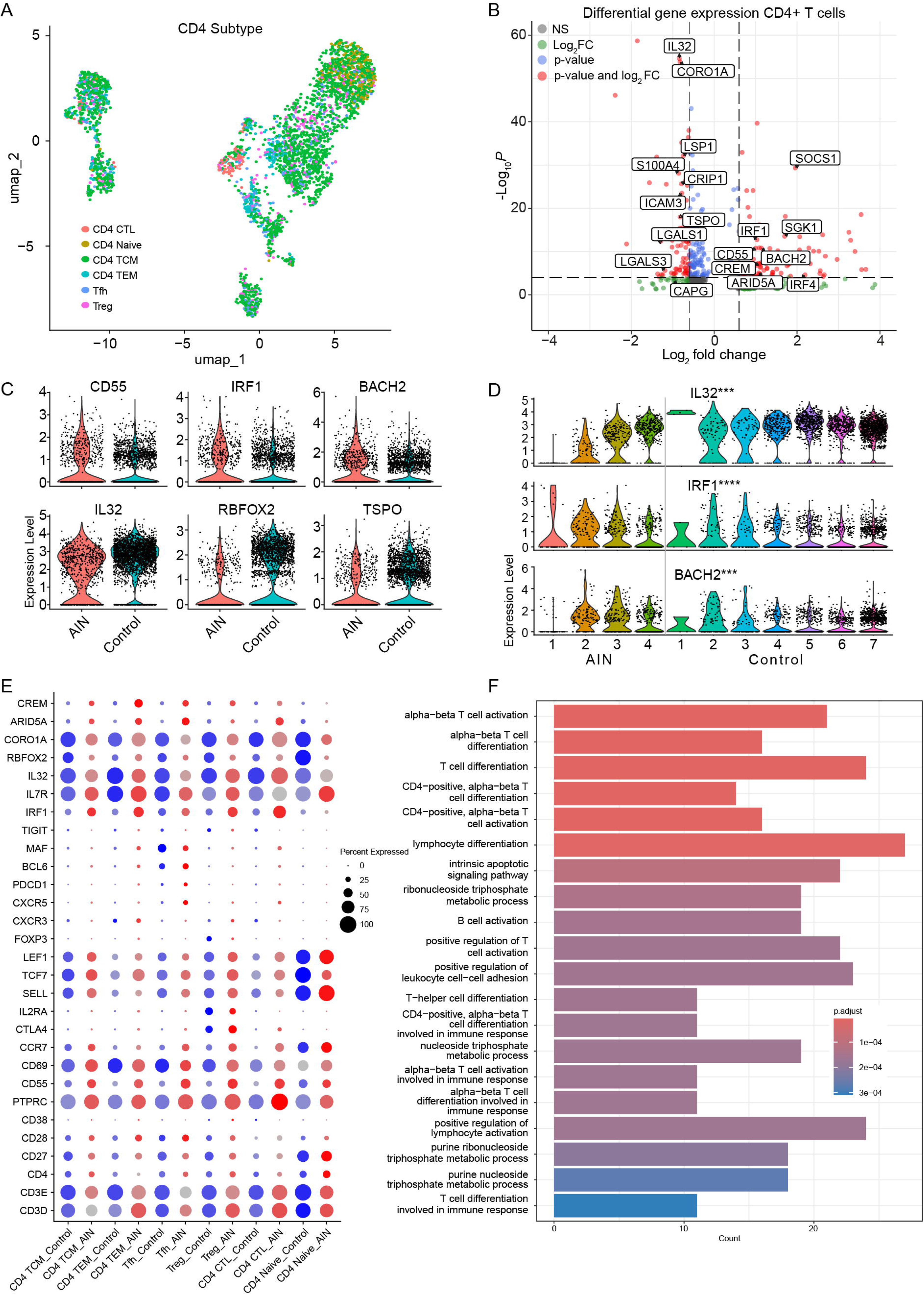
A) Uniform manifold approximation and projection (UMAP) of CD4+ T cells in AIN (n=4) and healthy controls (n=7) showing clustering of different cell populations (each population is indicated by a distinct color). B) Volcano plot shows differential gene expression in CD4+ T cells between AIN patients and healthy controls. The dotted horizontal line shows the p-value cut off and vertical lines show log2-fold change difference in expression. C-D) Representative examples of some differentially expressed genes between groups and at individual sample levels. E) Dot plot shows differential expression of selected T cell markers in the CD4+ T cell cohort. F) Bar plot shows top relevant pathways with gene count differentiating AIN patients and healthy controls. [AIN: neurofascin 155 mediated autoimmune nodopathy, HC: healthy control, **** p < 0.001, *** p < 0.005, ** p < 0.01, * p < 0.05]

### Differences in naïve B cells and CD4+ T cells are specific to NF155-AIN

To confirm whether the noted differences in naïve B cells and CD4+ T cells in NF155-AIN patients are specific to NF155-AIN, we examined transcriptomics profiles from three patients with CIDP, another acquired demyelinating disease of the peripheral nervous system, but without a known autoantibody associated with disease pathomechanism. Some of the key differences at the transcriptomics level of naïve B cells and CD4+ T cells appeared to be specific to NF155-AIN (**Fig. 6, Fig. S6, S7**), and overrepresentation analysis confirmed impaired differentiation and activation of B cells and CD4+ T cells, along with impaired B cell – CD4+ T cell interaction in NF155-AIN compared to CIDP. These findings are important because they suggest that NF155-AIN has a distinct underlying pathomechanism compared to CIDP, despite both being autoimmune diseases affecting peripheral nerves with altered immune environments.

**Figure 6.**
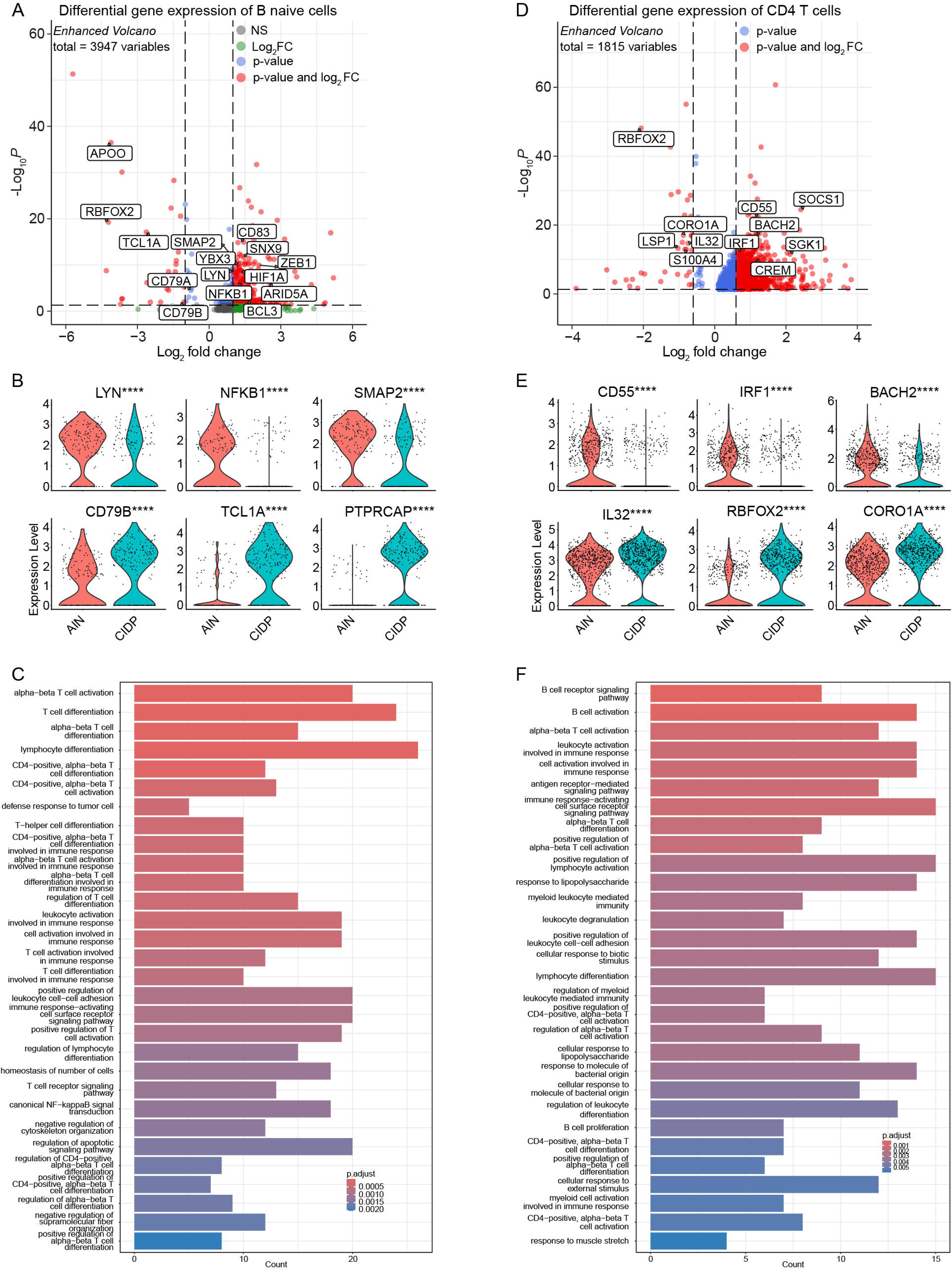
A) Volcano plot shows selected gene expression in naïve B cells between AIN and CIDP patients, and B) violin plot shows expression of some selected genes between groups in B naïve cells. C) Bar plot shows pathways impaired in naïve B cells between AIN and CIDP. D) Volcano plot shows selected gene expression in CD4+ T cells between AIN and CIDP patients, and E) violin plot shows expression of differentially expressed representative genes. F) Bar plot shows pathways altered in CD4+ T cells in AIN compared to CIDP. [AIN: neurofascin 155 mediated autoimmune nodopathy, CIDP: chronic inflammatory demyelinating polyneuropathy, **** p < 0.001, *** p < 0.005, ** p < 0.01, * p < 0.05]

### Impaired cell-cell communication in NF155-AIN

While examining the different compartments of immune cells provides a detailed understanding of individual compartments, the immune system is complex, and as explained above, interactions between different subtypes of immune cells are critical to maintain normal immune function. We leveraged the CellChat platform, which provides a comprehensive examination of ligand-receptor interactions, to examine the global differences in cell-cell communications in NF155-AIN and compared them with HC and CIDP. While the interaction number between NF155-AIN vs. HC was comparable, interaction strength was reduced in NF155-AIN, and several pathways were altered in NF155-AIN (**Fig. 7A, 7C**). Distinct differences in incoming and outgoing signals were noted in naïve B and Tfh cells in NF155-AIN compared to HC and CIDP (**Fig. 7B**). Details of incoming and outgoing signals between different immune cells (**Fig. 7C**), B cells and T cells- and signaling received by B cells from T cells are shown in **Fig. 7C-E**. Of interest are interactions mediated by ICAM, SEMA4, SELL, and SELPLG, which are genes involved in immune cell adhesion, migration, and costimulatory signaling.^62–66^ Finally, we examined the cell-cell communication among the altered pathways to ensure that they are specific to NF155-AIN, and among the noteworthy pathways, ADGRE and CypA were upregulated in NF155-AIN; and SELPLG was downregulated in NF155-AIN (**Fig. 7F, 7G, Fig. S8**). These findings suggest that the B naïve population in NF155-AIN is distinct, and abnormal ligand-receptor interactions between B naïve and Tfh cells may have played a role in B cell development.

**Figure 7.**
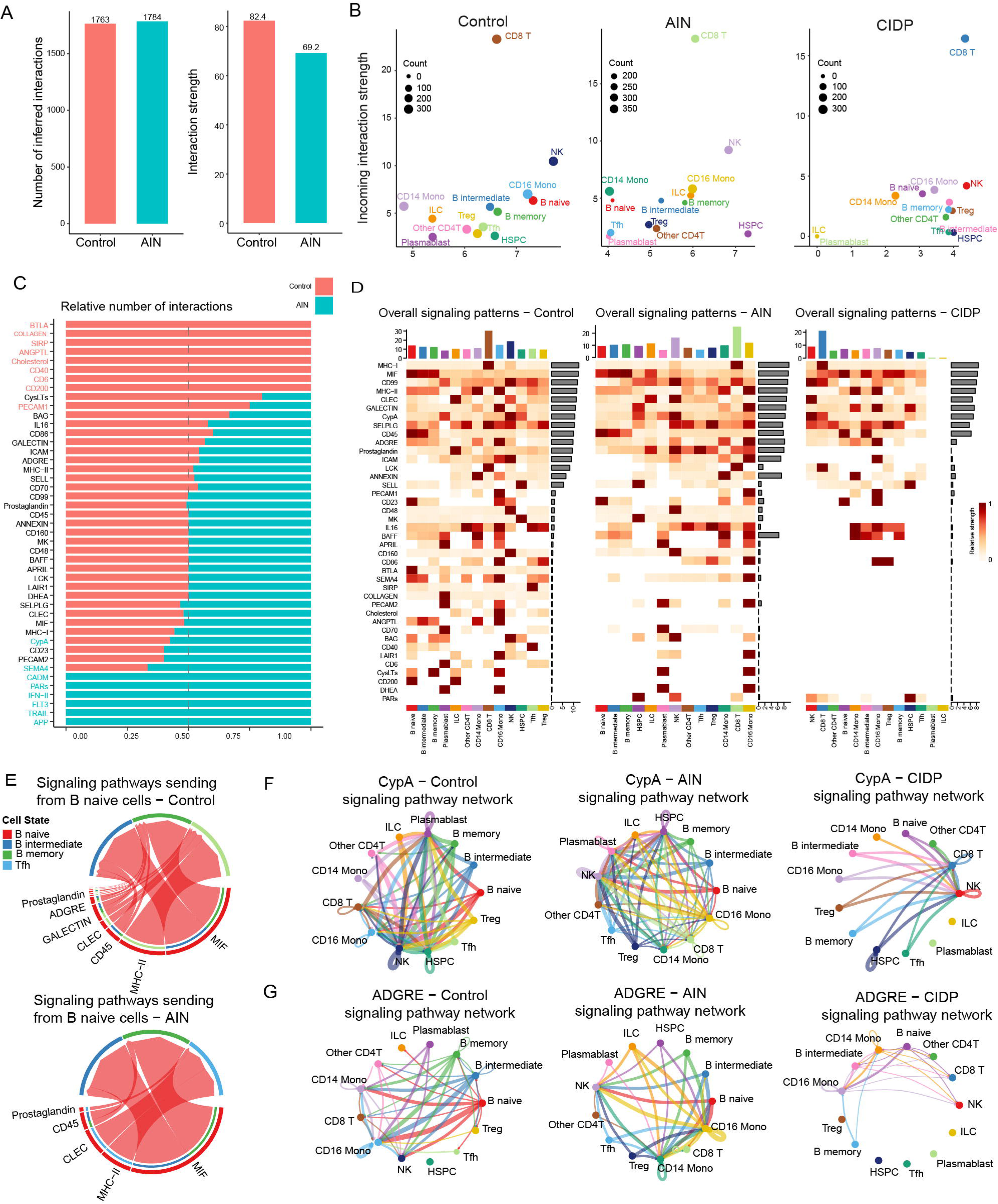
A) Bar plots show total number of possible interactions and strength of interactions between AIN patients and healthy controls. B) Overall signaling pattern from different cell types between healthy controls and AIN patients showing a distinct B naïve population in AIN patients, C) Bar plot shows significant signaling pathways ranked based on their differences of overall information flow, calculated by summarizing all communication probabilities in a given inferred network. Those colored orange and teal are more enriched in healthy controls and AIN patients, respectively. D) Overall signaling pattern between different types of cells in AIN, healthy control, and CIDP. E) Signaling pathways sent from B naïve cells in AIN and healthy controls. F-G) Circle plots of inferred CypA and ADGRE network in healthy controls, AIN, and CIDP patients. [AIN: neurofascin 155 mediated autoimmune nodopathy, HC: healthy control, CIDP: chronic inflammatory demyelinating polyneuropathy]

### Similarities in pathogenesis between NF155-AIN and MuSK+ MG, another IgG4-mediated disease

Finally, we examined the B naïve cell population in MuSK+ MG and HC to determine if there are any commonalities in the B naïve cell population between MuSK+ MG and NF155-AIN. We noted low expression of CD79B, TCL1A, and SELL in patients with MuSK+ MG with elevated CD83, HIF1A, and YBX3, similar to NF155-AIN (**Fig. 8B, C**). Restricted usage of IGHV3 genes was observed in patients with NF155-AIN, consistent with the restricted IGHV3 region usage previously reported in MuSK MG (**Fig. S3**).^16^ Furthermore, a recent report observed a lower expression of CD19+ in MuSK+MG, as in NF155-AIN, which could be linked to short-lived plasmablasts.^67^ These findings suggest a potentially similar pathomechanism in IgG4 autoantibody-mediated diseases.

**Figure 8.**
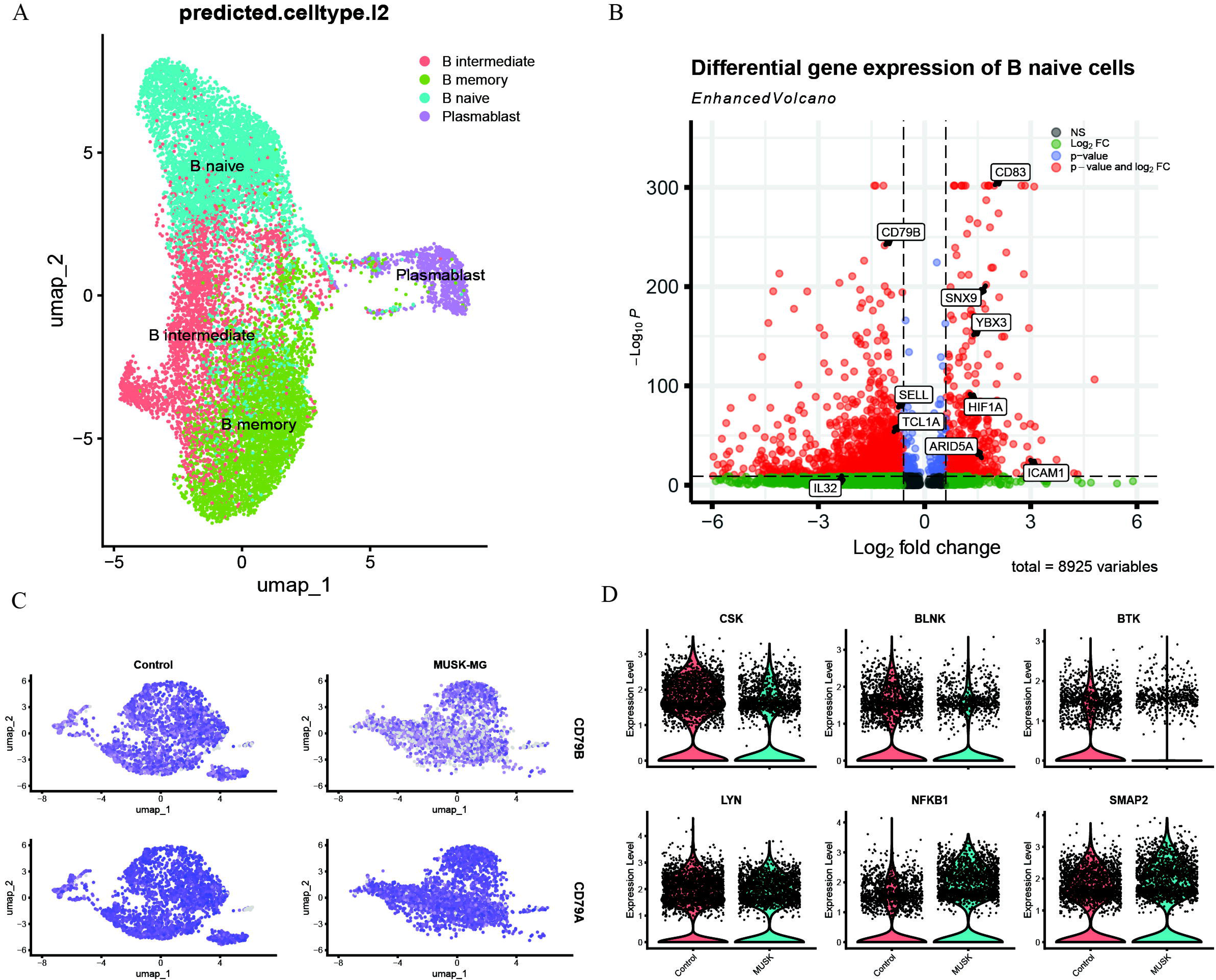
A) Uniform manifold approximation and projection (UMAP) of B cells in MuSK+ MG (n=3) and healthy controls (n=7) showing clustering of different cell populations (each population is indicated by a distinct color). B) Volcano plot shows differential gene expression in B naive cells between MuSK+ MG patients and healthy controls. Vertical lines show log2-fold change difference in expression. C) Feature plot showing expression of CD79A and CD79B in B naïve cells between MuSK+ MG and healthy controls. D) Violin plot showing expression of some of the relevant genes in B naïve cells between MuSK+ve MG and healthy controls. [MuSK-MG: Myasthenia gravis with autoantibody against muscle-specific tyrosine kinase, HC: healthy control]

## Discussion

Humoral immunity plays an important role in the pathogenesis of NF155-AIN, and autoantibodies against NF155 have been shown to be pathogenic in both *in vitro* and *in vivo* experiments.^5,7,68–71^ Our results showed an early B cell tolerance defect in NF155-AIN, with affected naïve B cell development, which is potentially mediated by impaired BCR signaling and regulation by CD4+ T cells. Furthermore, these differences appear specific to NF155-AIN when compared to CIDP—another demyelinating disease of the peripheral nerves that lacks known autoantibodies—and may reflect shared pathomechanisms with other IgG4 autoantibody-mediated diseases, such as MuSK+ MG.

B cell tolerance checkpoints are dysfunctional in many autoimmune diseases.^14,16,18–20,72^ Abnormal BCR/Toll-like receptor (TLR) plays a critical role in central B cell tolerance.^16^ Genome-wide association studies (GWAS) have previously shown that PTPN22 polymorphism is the second most important genetic risk factor after MHC in the development of autoimmune diseases, including rheumatoid arthritis, systemic lupus erythematosus (SLE), and type 1 diabetes. Moreover, a missense mutation in PTPN22 limits interaction with CSK tyrosine kinase and decreases TLR and BCR signaling.^73^ We noted abnormal BCR signaling in NF155-AIN that was independent of PTPN22 and potentially stemmed from low CD79B expression.^74^ Furthermore, it appears that pathways related to bis-phosphorylation of immunoreceptor tyrosine-based activation motif (ITAM), which recruits SYK and the downstream pathways involving BLNK and BTK, were more involved compared to recruitment of Lyn activation following mono-phosphorylation of ITAM. We also noted low expression of CSK, which is another C-terminal Src tyrosine kinase. Reduced CSK expression may therefore lead to a decreased threshold for Lyn activation.^73–77^

We observed upregulated intrinsic anti-apoptotic pathways in patients with NF155-AIN, including high expression of HIF1A, BCL2, and BCL3. HIF1A can be activated either by hypoxia or BCR signaling and plays an important role in the generation and quality of humoral immune response. HIF1A is usually more active in pro-B and pre-B cells and less active in immature B cells. The activation of HIF1A may lead to lower surface BCR and reduced CD19 expression, both noted in NF155-AIN.^78^ Furthermore, HIF1A may also rewire B cell metabolism and alter immunoglobulin production.^79^ Besides its apoptotic role, enforced expression of BCL2 has already been implicated in abrogating tolerance.^80^ Similarly, BCL3 is a pro-survival gene that also regulates NF-kB activation and has been implicated in B cell malignancies and other cancers.^81,82^ YBX3 is also an inhibitor of apoptosis, which supports cell survival through the PI3K/AKT pathway.^83^

Apart from the impaired BCR signaling and pro-survival genes, we also noted abnormal intracellular transport, with upregulated SMAP2 and SNX9, both associated with clathrin-mediated endocytosis of membrane compartments, in NF155-AIN. Previously, SNX9 downregulation has been reported in chronic inflammatory conditions but the role of SMAP2 in autoimmunity remains largely unexplored.^84^ Possibly, these proteins were upregulated in the context of low BCR expression as an attempt to normalize BCR internalization, which is dependent on clathrin-mediated endocytosis, to restore BCR signaling.^85^

There are other mechanisms involved in B cell tolerance. For example, defective receptor editing via genetic mutations in the recombinase-activating genes (RAGs) could also result in a highly autoreactive repertoire.^86–88^ Furthermore, alteration in BCR and TLR pathways including adenosine deaminase (ADA), WAS, MYD88, IRAK 4, transmembrane activator and CAML Interactor (TACI), and activation induced cytidine deaminase (AID) can affect central B cell tolerance.^40,89,90^ While our focus was an unbiased analysis of the naïve B cell and CD4+ T cell population, we also examined the expression of these particular genes in B cells and CD4+ T cells. WAS was downregulated in naïve B cells in NF155-AIN, compared to controls, but the difference was less striking compared to CIDP. We did not notice any major role of these genes in the development of dysfunctional early B cell tolerance in NF155-AIN, and overall expression of some of these genes, along with some key TLR genes, was low across the groups likely as a limitation of the single cell transcriptomics platform, making a meaningful statistical analysis not feasible.^40,91^

It is already known that regulatory T cells play a major role in peripheral B cell tolerance, and autoimmune regulator (AIRE) deficient patients display specific defects in the peripheral B cell tolerance checkpoint caused by a failure of AIRE mediated T cell/regulatory T cell (Treg) selection.^41^ Decreased Treg frequencies and/or impaired Treg suppressive function have been found in patients with ADA, CD40L, DOCK8, MHC class II-, and WAS-deficiency.^16^ We noted low interaction of CD40, and altered MHC pathway in NF155-AIN (**Fig. S8**).^92,93^ Furthermore, we also noted impaired CD4+ T cell development and homeostasis, and abnormal T regulatory cell function and regulation may affect peripheral tolerance.^55,94^

Among the different pathways, we also noted low SELL expression and abnormal SELPLG mediated interaction in NF155-AIN. SELL is critical for leukocyte migration, and previous studies have reported overlapping interaction between SELL^95^ and ICAM-1 pathways, and we also noted abnormal ICAM-1 pathway in NF155-AIN.^63^ However, the impact of low SELL and abnormal ICAM-1 pathway in B cell development, and whether it is linked to low BTK and WAS is not clear. Similarly, we noted overexpression of ADGRE and CD55. Previous work has suggested ADGRE5 is upregulated in T cells of patients treated with immune-checkpoint inhibitor, in the STAT5-IL-32 dependent pathway, but the exact role of ADGRE upregulation in CD4+ T cells in the context of NF155-AIN remains undefined.^96^ Similarly, CD55 overexpression inhibits complement activation, and previous work on SLE has shown decreased CD55 expression in lymphocytes in systemic lupus erythematosus and bullous pemphigoid patients,^97–99^ but why they are overexpressed in NF155-AIN needs further exploration, as the complement system is not considered to have a major role in NF155-AIN. Interestingly, CD97, a member of the EGF-TM7 protein family, is upregulated in activated T cells and binds CD55. Blocking of CD55-CD97 interaction has been shown to inhibit the proliferation of T cells and interferon γ secretion.^100^

While we observed several interesting findings, there are limitations to this study. The major limitation is sample size, which is unsurprising given the rarity of NF155-AIN. However, we have used robust statistical methods to address this limitation and validated our findings by comparing them with other relevant diseases, thereby reinforcing the reliability of our observations. As patients with NF155-AIN were treated with IVIg, we have used CIDP patients who were also treated with IVIg to limit the potential confounding from underlying therapy, but we could not adjust for the dose and time since the last IVIg administration. There are also standard technical limitations with single-cell analysis, and further confirmatory studies are warranted.^101^ Our analysis has primarily focused on naïve B cells and CD4 T cells; however, further investigation of additional immune cell compartments is warranted, including ongoing studies of BCR and TCR repertoires. In NF155-AIN, IgG4 autoantibodies are produced by plasmablasts and plasma cells, but their low frequency in peripheral blood limited our ability to perform in-depth transcriptomic analysis combined with surface phenotyping. Nonetheless, ongoing efforts are aimed at more precise immunophenotyping of these cells, which may also reveal novel therapeutic targets.

Despite these limitations, overall, our analysis confirmed a breach in B cell tolerance checkpoints in patients with NF155-AIN, which is potentially mediated by impaired BCR signaling and lack of regulation from CD4+ T cells. Our single-cell analysis confirmed some previously reported findings in NF155-AIN, reinforcing the validity of our findings. Furthermore, the unbiased transcriptomics study identified several new aspects of NF155-AIN. While confirming the implications of each of these new findings was beyond the scope of the present work, these findings will guide future direction of research in NF155-AIN, a rare but potentially debilitating disease, and may eventually lead to new therapeutic discoveries to improve patient care.

## Data availability

Relevant single-cell transcriptomics data will be deposited in Gene Omnibus upon the publication of the manuscript.

## Acknowledgements

The authors acknowledge the support of the patients who participated in this research by donating blood samples, and agreed to share their de-identified clinical information, which made this research possible. We also acknowledge the help in single-cell experiments and analysis from Guilin Wang, Arun Chavan, and YCGA, and Beata Filipek for her feedback on the manuscript.

## Funding

This research was supported by the NIH award 1R01NS132860-01A1.

KOC is supported by NIAID under award R01AI114780.

RJN is supported by the National Institutes of Health funded Rare Diseases Clinical Research Network (NIH-RDCRN) under award number U54NS115054 (MGNet).

LMA. was supported by a personal Juan Rodés grant JR21/00060 and LQ by a personal clinical intensification INT23/00066 and project PI22/00387 by Fondo de Investigaciones Sanitarias (FIS), Instituto de Salud Carlos III (Spain).

## Competing interests

BR has served as a consultant/advisor for Alexion (now part of AstraZeneca), Takeda, Sanofi, and argenx. Additionally, BR has received research support from the Martin Shubik Fund for IBM at Yale University, NIH, Abcuro Pharmaceuticals, argenx, Immunovant, and Takeda. BR has small stocks in Cabaletta Bio and Pfizer.

KCO is an equity shareholder of Cabaletta Bio; serves on advisory boards for Roche, Merck (EMD Serono), Neurocrine Biosciences, and Seismic Therapeutic; and has received research support from Viela Bio, (now Horizon Therapeutics/Amgen), argenx, and Seismic Therapeutic.

RJN has received research support from the NIH, Genentech, Alexion (Astra Zeneca), argenx, Annexon Biosciences, Ra Pharmaceuticals (now UCB), Myasthenia Gravis Foundation of America, Momenta (now Janssen), Immunovant, Grifols, and Viela Bio (Horizon Therapeutics, now Amgen). RJN has also served as a consultant/advisor for Alexion (Astra Zeneca), argenx, Cabaletta Bio, CSL Behring, Grifols, Ra Pharmaceuticals (now UCB Pharma), Immunovant, Momenta (now Janssen), Viela Bio (Horizon Therapeutics, now Amgen).

ALH has served as a consultant for argenx, Alexion, UCB, Genentech, EMD Serono.

AO and CC did not have any conflict of interest when working on this project. AO is now employed by AbbVie Inc. CC is now employed by Merck Serono.

JC, KGK, SO, FNCK, DJ, GM, SD, and CL have nothing to disclose. Hospital de la Santa Creu i Sant Pau receives payment for autoimmune nodopathies antibody testing.

LQ received research grants from Instituto de Salud Carlos III – Ministry of Economy and Innovation (Spain), CIBERER, Fundació La Marató, GBS-CIDP Foundation International and ArgenX. LQ received speaker or expert testimony honoraria from CSL Behring, Novartis, Sanofi-Genzyme, Merck, Annexon, Alnylam, ArgenX, Dianthus, Biocryst, Montis, LFB, Avilar Therapeutics, Lycia Therapeutics, Nuvig Therapeutics and Takeda. LQ serves at Clinical Trial Steering Committees for Sanofi Genzyme, ArgenX and Takeda and is Principal Investigator for UCB’s CIDP01 trial and Sanofi’s Mobilize and Vitalize trials.

---

**Supplementary Figure S1**. Confirming proper integration of samples showing uniform manifold approximation and projection (UMAP) of peripheral blood mononuclear cells (PBMCs) A) between cells from healthy controls and AIN patients, and B) cells from different samples, for Figure 3A. UMAP of B cells only from C) cells from healthy controls and AIN patients, and D) cells from individual samples. UMAP of B cells only from E) healthy controls and CIDP patients, and F) cells from individual samples. UMAP of B cells only from G) cells from MuSK+ MG and healthy controls, and H) cells from individual samples.

[AIN: NF155 mediated AIN, MuSK-MG: Myasthenia gravis with autoantibody against muscle-specific tyrosine kinase]

**Supplementary Figure S2.** Top 5 gene expression between clusters of B cells from NF155-AIN and healthy controls.

**Supplementary Figure S3**. Uniform manifold approximation and projection (UMAP) of B naïve cells from A) AIN patients and healthy controls, B) cells from individual samples. C) Differential gene expression between AIN and healthy control for top features in each cluster of naïve B cells. D) Feature plot shows expression of some representative genes between AIN patients and healthy controls.

**Supplementary Figure S4.** A) Volcano plot shows differential gene expression in intermediate B cells between AIN patients and healthy controls. The dotted horizontal line shows the p-value cut off and vertical lines show log2-fold change difference in gene expression. B) Bar plot shows top relevant pathways with gene count differentiating AIN patients and healthy controls. C-D) Violin plots show expression of some important genes between groups and between individual samples.

**Supplementary Figure S5.** A) Volcano plot shows differential gene expression in memory B cells between AIN patients and healthy controls. The dotted horizontal line shows the p-value cut off and vertical lines show log2-fold change in gene expression. B) Bar plot shows top relevant pathways with gene count differentiating AIN patients and healthy controls. C-D) Violin plots show expression of some important genes between groups and between individual samples.

**Supplementary Figure S6 A, B)** Feature plots and violin plots show expression of selected genes of interest in naïve B cells between AIN and CIDP patients. **C, D**) Heatmap showing expression of different genes between AIN and control, and CIDP and control.

**Supplementary Figure S7.** Feature plot shows expression of selected genes of interest in CD4+ T cells between AIN and CIDP patients, and expression of some selected genes at the individual level are shown in violin plots.

**Supplementary Figure S8.** A-C) Chord diagrams show connections of different cell types in some selected pathways and related gene expression in NF155-AIN patients compared to healthy controls. D-E) Expression of ADGRE and CypA pathway related genes in healthy controls (orange) and NF155-AIN patients (green).

