## Supplementary Tables for "B Cell Tolerance and BCR Signaling Dysregulation in NF155-Mediated Autoimmune Nodopathies"

**Supplementary Table 1.**

| **Subjects for the B cell tolerance checkpoint assessment study** | | | | | | | | |
| --- | --- | --- | --- | --- | --- | --- | --- | --- |
| Sample | Age, years | Sex | Clinical Presentation | Electrodiagnostic Information | anti-NF155 | Treatment received | Response to Rituximab | Other clinical comorbidities at the time of sample collection |
| AIN-1 | 56 | F | Proximal>distal motor weakness, distal sensory loss, areflexic | Demyelinating neuropathy | IgG | IVIG | N/A | Gastro-esophageal reflux disease, Migraine, Breast Cancer (remote) |
| AIN-2 | 61 | M | Distal motor weakness, distal sensory deficit, areflexic, ataxia | Demyelinating neuropathy | IgG4 | IVIg, glucocorticoids, PLEX | Yes | hypertension |
| AIN-3 | 43 | M | Pure sensory symptoms, Areflexic | Demyelinating neuropathy | IgG4 | glucocorticoids | Partial | hypertension |
| HC-1 | 30 | F | NA | NA | NA | NA | NA | None |
| HC-2 | 47 | M | NA | NA | NA | NA | NA | None |
| HC-3 | 53 | M | NA | NA | NA | NA | NA | None |
| HC-4 | 28 | F | NA | NA | NA | NA | NA | None |
| **Subjects for single-cell transcriptomics study (AIN)** | | | | | | | | |
| **AIN samples** | | | | | | | | |
| AIN-1 | 61 | M | Distal motor weakness, distal sensory deficit, areflexic, ataxia | Demyelinating neuropathy | IgG4 | IVIg, glucocorticoids, PLEX | Yes | hypertension |
| AIN-2 | 43 | M | Pure sensory symptoms, Areflexic | Demyelinating neuropathy | IgG4 | glucocorticoids | Partial | hypertension |
| AIN-3 | 53 | M |  |  | IgG4 |  |  |  |
| AIN-4 | 26 | F | Proximal and distal motor weakness, facial weakness, distal sensory deficit, areflexic, ataxia | Demyelinating neuropathy | IgG4 | IVIg | Partial | Polycystic ovarian syndrome, non-secreting pituitary adenoma |
| **Chronic inflammatory demyelinating polyneuropathy (CIDP) samples** | | | | | | | | |
| CIDP-2 | 41 | M |  | Demyelinating neuropathy |  | IVIg | N/A | None |
| CIDP-3 | 67 | F |  | Demyelinating neuropathy |  | IVIg | N/A | Hypertension |
| CIDP-1 | 56 | F |  | Demyelinating neuropathy |  | IVIg | N/A | Remote history of breast cancer |
| **MuSK+ MG samples** | | | | | | | | |
| MuSK-1 | 43 | F | Ptosis of right eyelid, diplopia, then developed bulbar symptoms | NA |  | Rituximab over 12 months ago | Yes | None |
| MuSK-2 | 65 | F | Bulbar symptoms | NA |  | Rituximab over 12 months ago | Yes | None |
| MuSK-3 | 59 | F | Bulbar symptoms | NA |  | Plasmapheresis | NA | hypertension |
| **Healthy controls (HC)** | | | | | | | | |
| HC-1 | 30 | M |  |  |  |  |  | None |
| HC-2 | 39 | M |  |  |  |  |  | None |
| HC-3 | 55 | M |  |  |  |  |  | None |
| HC-4 | 59 | M |  |  |  |  |  | hypertension |
| HC-5 | 72 | M |  |  |  |  |  | hypertension |
| HC-6 | 37 | F |  |  |  |  |  | None |
| HC-7 | 50 | F |  |  |  |  |  | hypertension |

**Supplementary Table 1**: Demographic details of patients with AIN and healthy controls included in this study. None of the patients received rituximab prior to sample collection.

[AIN: autoimmune nodopathy, CIDP- Chronic inflammatory demyelinating polyneuropathy, HC: healthy control, MuSK+ MG: Muscle specific tyrosine kinase autoantibody positive myasthenia gravis (MG)]

**Table S2**. Sequence and reactivity tables for NF155-AIN and HD subjects

Sequence and reactivity tables for new emigrant B cells from NF155-AIN1

|  | HEAVY | | | | | | |  | LIGHT | | | | |  | REACTIVITY |
| --- | --- | --- | --- | --- | --- | --- | --- | --- | --- | --- | --- | --- | --- | --- | --- |
| rIgG | VH | D | RF | JH | CDR3(aa) | Length | Mutations |  | Vk | Jk | CDR3(aa) | Length | Mutations |  | Poly |
| aF11 | 1-2 | 4-17 | 2 | 3 | ARDLRLDDYGDYGGFDI | 17 | 0 | kappa | 3-20 | 1 | QQYGSSVT | 8 | 0 |  | - |
| aF6 | 3-30-3 | 4-17 | 3 | 2 | ARDPMTTVTTIGWYFDL | 17 | 0 | kappa | 1-39 | 5 | QQSYSTPIT | 9 | 0 |  | - |
| aF7 | 4-61 | 4-17 | 3 | 5 | ARMTTVTIA | 9 | 0 | kappa | 3-11 | 1 | QQRSNWPPSWT | 11 | 0 |  | + |
| aF9 | 3-23 | 3-22 | 2 | 4 | AKPRPPSSGYSYFDY | 15 | 0 | kappa | 1-5 | 3 | QQYNSYPFT | 9 | 1 m CDR3 |  | - |
| aG3 | 3-23 | 5-12 | 3 | 3 | AKDRGRGSGYEGAFDI | 16 | 0 | kappa | 1-33 | 3 | QQYDNLPFT | 9 | 0 |  | - |
| aG9 | 1-3 | 3-22 | 2 | 4 | ARARGSSGYYYGWTLDY | 17 | 0 | kappa | 2-28 | 4 | MQALQTPT | 8 | 0 |  | - |
| aH11 | 4-38-2 | 3-10 | 2 | 4 | ARHWAYYYGSGHHFDY | 16 | 0 | kappa | 3-20 | 4 | QQYGSSPLT | 9 | 0 |  | + |
| aH5 | 3-74 | 3-22 | 2 | 4 | ARAPKRGEYYYDSSGYSPDY | 20 | 0 | kappa | 1-17 | 2 | LQHNSFPYT | 9 | 1 m CDR3 |  | - |
| rIgG | VH | D | RF | JH | CDR3(aa) | Length | Mutations |  | Vλ | Jλ | CDR3(aa) | Length | Mutations |  | Poly |
| aF1 | 3-30 | 2-15 | 3 | 4 | ARGIVVVAATTPGGFDY | 17 | 0 | lam | 3-25 | 2 | QSADSSGTYYVV | 12 | 0 |  | - |
| aF2 | 3-15 | 6-19 | 1 | 4 | TTSRRSGSDDYFDY | 14 | 0 | lam | 3-1 | 2 | QAWDSSSVV | 9 | 0 |  | - |
| aG4 | 3-21 | 3-22 | 2 | 4 | ARSPTYYYDSSGHYYADY | 18 | 1 m CDR1 | lam | 1-47 | 3 | AAWDDSLSGRV | 11 | 0 |  | + |

RF: reading frame	-: non-reactive	+: reactive	FR: framework	CDR: complementarity-determining region	m: missense mutation

Sequence and reactivity tables for mature naïve B cells from NF155-AIN1

|  | HEAVY | | | | | | |  | LIGHT | | | | |  | REACTIVITY |
| --- | --- | --- | --- | --- | --- | --- | --- | --- | --- | --- | --- | --- | --- | --- | --- |
| rIgG | VH | D | RF | JH | CDR3(aa) | Length | Mutations |  | Vk | Jk | CDR3(aa) | Length | Mutations |  | Poly |
| aB2 | 1-3 | 1-26 | 1 | 4 | ARAVGATSGY | 10 | 1 m FR3 |  | 4-1 | 2 | QQYYSTPYT | 9 | 0 |  | - |
| aC11 | 1-69 | 6-13 | 2 | 6 | ARAGAAAGTQYYYYMDV | 17 | 0 |  | 3-20 | 4 | QQYGSSPNT | 9 | 0 |  | - |
| aC2 | 1-69 | 2-15 | 2 | 5 | ARDGGSSWYDY | 11 | 0 |  | 4-1 | 1 | QQYYSTPWT | 9 | 0 |  | + |
| aC9 | 4-59 | 5-18 | 1 | 6 | ARVPWGDTAMDYYYMDV | 17 | 0 |  | 2-28 | 1 | MQALQTPWT | 9 | 0 |  | - |
| aD2 | 4-61 | 1-1 | 2 | 5 | ARGKVHLGET | 10 | 0 |  | 1-39 | 4 | QQSYSTPLT | 9 | 0 |  | - |
| aD8 | 3-73 | 4-17 | 3 | 1 | TRHGSTVT | 8 | 0 |  | 4-1 | 1 | QQYYSTPWT | 9 | 0 |  | - |
| rIgG | VH | D | RF | JH | CDR3(aa) | Length | Mutations |  | Vλ | Jλ | CDR3(aa) | Length | Mutations |  | Poly |
| aB3 | 1-46 | 3-16 | 2 | 4 | AREPPGWKNRYYFDY | 15 | 0 |  | 2-23 | 2 | CSYAGSSTFPVV | 12 | 0 |  | + |
| aB6 | 4-34 | 2-21 | 2 | 4 | ARGRGYSLLY | 10 | 0 |  | 2-23 | 1 | CSYAGSSTSLNYV | 13 | 0 |  | + |
| aC10 | 3-53 | 5-24 | 3 | 4 | AREGEGYSFDY | 11 | 0 |  | 3-21 | 3 | QVWDSSSDHPRV | 12 | 0 |  | - |
| aC12 | 1-18 | 4-17 | 2 | 4 | ARVSLPIVIGDYGPFDY | 17 | 0 |  | 1-51 | 1 | GTWDSSLSAYV | 11 | 0 |  | - |
| aC3 | 1-18 | 3-3 | 3 | 3 | AREITIFGVVIKGAFDI | 17 | 0 |  | 2-11 | 2 | CSYAGSYTLV | 10 | 0 |  | - |
| aC4 | 4-59 | 3-16 | 2 | 4 | ARHHYDYVWGSYRTPYYFDY | 20 | 0 |  | 8-61 | 3 | VLYMGSGIWV | 10 | 0 |  | + |
| aC6 | 1-18 | 2-8 | 2 | 6 | ARGYSHCTGGVCLPGHYYMDV | 21 | 1 m FR3 |  | 1-36 | 3 | AAWDDSLNARV | 11 | 0 |  | - |
| aD10 | 3-30 | 6-19 | 2 | 4 | ARDGPVAGTMILYYFDY | 17 | 0 |  | 3-21 | 2 | QVWDSSSDHPVV | 12 | 1 m CDR2 |  | - |

RF: reading frame	-: non-reactive	+: reactive	FR: framework	CDR: complementarity-determining region	m: missense mutation

Sequence and reactivity tables for new emigrant B cells from NF155-AIN2

|  | HEAVY | | | | | | |  | LIGHT | | | | |  | REACTIVITY |
| --- | --- | --- | --- | --- | --- | --- | --- | --- | --- | --- | --- | --- | --- | --- | --- |
| rIgG | VH | D | RF | JH | CDR3(aa) | Length | Mutations |  | Vk | Jk | CDR3(aa) | Length | Mutations |  | Poly |
| aE11P1 | 4-34 | 3-22 | 2 | 4 | ARLLGDSSGYYEFDY | 15 | 0 |  | 1-17 | 1 | LQHNSYPRT | 9 | 0 |  | - |
| aF10P1 | 3-21 | 1-7 | 1 | 4 | ARGSITGTRERRPFDY | 16 | 0 |  | 3-11 | 1 | QQRSNWPPT | 9 | 0 |  | + |
| aF6HP1 | 4-31 | 6-25 | 3 | 4 | ARALPEHRFDY | 11 | 0 |  | 1-39 | 1 | QQSYSTPWT | 9 | 0 |  | - |
| aF8HP1 | 1-18 | 6-13 | 2 | 6 | ARGIAAAGTDYGMDV | 15 | 0 |  | 3-15 | 2 | QQYNNWPPYT | 10 | 0 |  | - |
| aG5HP1 | 3-33 | 2-21 | 2 | 4 | ARDPRAYCGGDCYLPLDY | 18 | 0 |  | 3-15 | 1 | QQYNNWPPWT | 10 | 0 |  | - |
| aE11P2 | 1-46 | 1-26 | 3 | 4 | ARDLDSGSFAWFDY | 14 | 0 |  | [1-39](https://www.imgt.org/IMGT_vquest/analysis#sequence1_alv) | [1](https://www.imgt.org/IMGT_vquest/analysis#sequence1_alj) | QQSYSTPRT | 9 | 0 |  | - |
| aF5 P2 | 3-7 | 3-22 | 3 | 3 | ARDGLVVITPNDAFDI | 16 | 0 |  | [1-39](https://www.imgt.org/IMGT_vquest/analysis#sequence19_alv) | [3](https://www.imgt.org/IMGT_vquest/analysis#sequence19_alj) | QQSYSTPLFT | 10 | 1 m FR3 |  | - |
| aH12P2 | 3-23 | 2-15 | 2 | 6 | AKRGSCSGGSCYGPDYYYYGMDV | 23 | 0 |  | [3-20](https://www.imgt.org/IMGT_vquest/analysis#sequence4_alv) | [2](https://www.imgt.org/IMGT_vquest/analysis#sequence4_alj) | QQYGSSPPGYT | 11 | 0 |  | + |
| aH3P2 | 3-15 | 6-13 | 2 | 4 | TTDPGAKPAEGIAAAGRHDY | 20 | 1 m FR3 |  | [1-5](https://www.imgt.org/IMGT_vquest/analysis#sequence13_alv) | [2](https://www.imgt.org/IMGT_vquest/analysis#sequence13_alj) | QQYNSYSYT | 9 | 0 |  | + |
| aE1 P3 | 3-9 | 6-19 | 1 | 5 | AKSRGISSGWYSYHFDP | 17 | 0 |  | 1-5 | 2 | QQYNSYSPRYT | 11 | 0 |  | - |
| aE7 P3 | 3-48 | 3-22 | 2 | 1 | ARDHLPADYYDSSGYYYRAEYFQH | 24 | 0 |  | 3-11 | 1 | QQRSNWPPWT | 10 | 0 |  | - |
| aG12P3 | 3-23 | 6-13 | 2 | 3 | AKDREAAAGTRGFGGGAFDI | 20 | 0 |  | 1-17 | 1 | LQHNSYPWT | 9 | 0 |  | + |
| aG1P3 | 3-39 | 3-3 | 3 | 3 | TRKPGDFGVVISAFDI | 16 | 0 |  | 1-6 | 1 | LQDYNYPRT | 9 | 0 |  | + |
| rIgG | VH | D | RF | JH | CDR3(aa) | Length | Mutations |  | Vλ | Jλ | CDR3(aa) | Length | Mutations |  | Poly |
| aE3HP1 | 3-30 | 3-22 | 2 | 2 | AKDGGDYYDSSGYTFWWYFDL | 21 | 1 m CDR3 |  | 1-40 | 2 | QSYDSSLSGSVV | 12 | 1 m FR3 |  | - |
| aE6HP1 | 1-18 | 2-15 | 2 | 4 | ARARSGGGSLDY | 12 | 0 |  | 3-21 | 2 | QVWDSSSDHPV | 11 | 0 |  | - |
| aG4HP1 | 3-53 | 4-17 | 3 | 4 | ARGTVTVAFDY | 11 | 0 |  | 2-8 | 2 | SSYAGSNNFVV | 11 | 0 |  | - |
| aG6 P2 | 1-69 | 1-69 | 2 | 4 | ARGNGASWHGDSFDY | 15 | 1 m FR2 |  | [1-47](https://www.imgt.org/IMGT_vquest/analysis#sequence11_alv) | [2](https://www.imgt.org/IMGT_vquest/analysis#sequence11_alj) | AAWDDSLSGPV | 11 | 0 |  | + |

RF: reading frame	-: non-reactive	+: reactive	FR: framework	CDR: complementarity-determining region	m: missense mutation

Sequence and reactivity tables for mature naïve B cells from NF155-AIN2

|  | HEAVY | | | | | | | |  | LIGHT | | | | |  | REACTIVITY | |
| --- | --- | --- | --- | --- | --- | --- | --- | --- | --- | --- | --- | --- | --- | --- | --- | --- | --- |
| rIgG | VH | D | RF | JH | CDR3(aa) | Length | Mutations |  | Vk | | Jk | CDR3(aa) | Length | Mutations |  | | Poly |
| aB11P1 | 1-18 | 1-26 | 1 | 4 | AREGWGIVGATNAADY | 16 | 0 |  | 2-40 | | 2 | MQRIEFPYT | 9 | 0 |  | | - |
| aD5HP1 | 4-34 | 6-19 | 2 | 4 | ARGNAPTVAAYFDY | 14 | 0 |  | 5-3 | | 1 | QQYNSSGT | 8 | 0 |  | | - |
| aD9P2 | 1-18 | 7-27 | 2 | 3 | ARDLGDAFDI | 10 | 0 |  | [3-11](https://www.imgt.org/IMGT_vquest/analysis#sequence16_alv) | | [4](https://www.imgt.org/IMGT_vquest/analysis#sequence16_alj) | QQRRST | 6 | 0 |  | | - |
| aA1P3 | 3-53 | 4-17 | 2 | 3 | ARAVVGYGDYDRLVAFDI | 18 | 0 |  | 2-28 | | 4 | MQALQTPLT | 9 | 0 |  | | - |
| aA4P3 | 4-59 | 2-2 | 2 | 3 | ARNGVNKKYCSSTSCYQMNSDAFDV | 25 | 0 |  | [1-33](https://www.imgt.org/IMGT_vquest/analysis#sequence2_alv) | | [4](https://www.imgt.org/IMGT_vquest/analysis#sequence2_alj) | QQYDNLPLT | 9 | 0 |  | | + |
| aB12P3 | 3-33 | 3-9 | 1 | 6 | ARDELRYFDWPPPAYYYGMDV | 21 | 0 |  | 2-28 | | 3 | MQALQTPLFT | 10 | 0 |  | | + |
| aC7P3 | 1-58 | 1-26 | 1 | 4 | AAGVGAIGY | 9 | 0 |  | 1-33 | | 4 | QQYDNLPRI | 9 | 0 |  | | + |
| aD12P3 | 3-48 | 2-21 | 3 | 6 | ARVTANYYYYGMDV | 14 | 0 |  | 2-28 | | 1 | MQALQTPRT | 9 | 0 |  | | - |
| rIgG | VH | D | RF | JH | CDR3(aa) | Length | Mutations |  | Vλ | | Jλ | CDR3(aa) | Length | Mutations |  | | Poly |
| aB12P1 | 3-74 | 5-12 | 3 | 3 | ARKRGYNGYGDDAFDI | 16 | 1 m CDR1, 1 m FR3 |  | 3-25 | | 3 | QSADSSGTYWV | 11 | 0 |  | | - |
| aC5HP1 | 3-7 | 6-19 | 2 | 4 | ARANQRIAVAGLPLDY | 16 | 0 |  | 1-40 | | 1 | QSYDSSLSGYV | 11 | 0 |  | | - |
| aC2 P3 | 3-33 | 6-19 | 1 | 5 | ARDGHPGWSSGWPENWFDP | 19 | 1 m FR2 |  | 2-11 | | 3 | CSYAGSYTWV | 10 | 0 |  | | - |
| aD2P3 | 3-23 | 1-26 | 1 | 3 | AKDLEGARRGVIVGATIDI | 19 | 0 |  | 3-21 | | 2 | QVWDSSSDHPDVV | 13 | 0 |  | | + |

RF: reading frame	-: non-reactive	+: reactive	FR: framework	CDR: complementarity-determining region	m: missense mutation

Sequence and reactivity tables for new emigrant B cells from NF155-AIN3

|  | HEAVY | | | | | | |  | LIGHT | | | | |  | REACTIVITY |
| --- | --- | --- | --- | --- | --- | --- | --- | --- | --- | --- | --- | --- | --- | --- | --- |
| rIgG | VH | D | RF | JH | CDR3(aa) | Length | Mutations |  | Vk | Jk | CDR3(aa) | Length | Mutations |  | Poly |
| aE6 | 4-34 |  | 3 | 4 | ARSFGVRGDLGY | 12 | 0 |  | 1-17 | 4 | LQHNSYPRT | 9 | 0 |  | - |
| aE9 | 1-2 | 3-22 | 2 | 4 | ARDGAYYDSSGYRQ | 14 | 0 |  | 1-17 | 1 | LQHNSYPRT | 9 | 0 |  | - |
| aF5 | 3-15 | 2-15 | 2 | 5 | TTELCSGGSCYSAVGKRPAR | 20 | 0 |  | 1-5 | 2 | QQYNSYPYT | 9 | 0 |  | - |
| aF6 | 3-11 | 3-22 | 2 | 6 | ARGHYDSSGYYYYYYYGMDV | 20 | 0 |  | 3-15 | 1 | QQYNNWPPWT | 10 | 0 |  | - |
| aG1 | 4-34 | 2-15 | 2 | 4 | ARGRRQKRVCSGGSCYYWGKEPLDY | 25 | 0 |  | 4-1 | 4 | QQYYSTPPT | 9 | 0 |  | + |
| aG10 | 4-61 | 6-6 | 2 | 6 | ARFGRIAARSGDYYYYYGMDV | 21 | 0 |  | 1-5 | 2 | QQYNSYYT | 8 | 0 |  | + |
| aG12-S1P1 | 4-59 | 6-6 | 1 | 6 | AREVSSSSRRLVYGMDV | 17 | 0 |  | 3-20 | 1 | QQYGSSPRT | 9 | 0 |  | - |
| aG12-S2P2 | 4-34 | 5-18 | 2 | 5 | ARGRRIQLWLRPTRVFDP | 18 | 0 |  | 4-1 | 1 | QQYYSTPQT | 9 | 0 |  | + |
| aG2 | 3-23 | 6-6 | 1 | 4 | AKVWWGCSSSSYFDY | 15 | 0 |  | 3-20 | 4 | QQYGSSPLT | 9 | 0 |  | + |
| rIgG | VH | D | RF | JH | CDR3(aa) | Length | Mutations |  | Vλ | Jλ | CDR3(aa) | Length | Mutations |  | Poly |
| aE2- s2p2 | 1-69 | 24*0 | 3 | 4 | ASTLSRDGYNYLSN | 14 | 0 |  | 2-14 | 2 | SSYTSSSTLVV | 11 | 1 CDR1 |  | + |
| aG12-p3 | 3-23 | 6-13 | 1 | 5 | ARSPSSSWYRWFDP | 14 | 0 |  | 2-8 | 3 | SSYAGSNNLV | 10 | 0 |  | - |
| aH4 | 3-23 | 3-10 | 2 | 6 | AKAYFSLSLLYGMDV | 15 | 0 |  | 6-57 | 2 | QSYDSSIYVV | 10 | 0 |  | + |

RF: reading frame	-: non-reactive	+: reactive	FR: framework	CDR: complementarity-determining region	m: missense mutation

Sequence and reactivity tables for mature naïve B cells from NF155-AIN3

|  | HEAVY | | | | | | |  | LIGHT | | | | |  | REACTIVITY |
| --- | --- | --- | --- | --- | --- | --- | --- | --- | --- | --- | --- | --- | --- | --- | --- |
| rIgG | VH | D | RF | JH | CDR3(aa) | Length | Mutations |  | Vk | Jk | CDR3(aa) | Length | Mutations |  | Poly |
| aB11-S2P2 | 3-48 | D1-7 | 2 | 5 | ARAAELELRFFWFDP | 15 | 0 |  | 1D-12 | 4 | QQANSFPLT | 9 | 0 |  | - |
| aB12- S1P2 | 1-3 | 4-17 | 3 | 4 | ARGIQPTVPSGD | 12 | 0 |  | 1-17 | 2 | LQHNSYPYT | 9 | 0 |  | - |
| aB2 s1p1 | 3-23 | 1-26 | 1 | 4 | AKCRRVGATTDPLDY | 15 | 0 |  | 1-33 | 1 | QQYDNLGA | 8 | 0 |  | + |
| aB3 s2P3 | 1-24 | 3-16 | 3 | 4 | ATVRIGGSYVNY | 12 | 0 |  | 3-11 | 1 | QQRSNWPRLWT | 11 | 0 |  | - |
| aB5 s2p2 | 3-23 | D1-7 | 1 | 4 | AKLSRGLTGTILYYFDC | 17 | 0 |  | 3-11 | 2 | QQRSNWPYT | 9 | 0 |  | + |
| aB7 s2p3 | 4-61 | 2-15 | 2 | 4 | ARGRGYCSGGSCYGPYYFDY | 20 | 0 |  | 3-11 | 4 | QQRSNWPLT | 9 | 0 |  | + |
| aC11 -s1p1 | 3-23 | 3-22 | 2 | 4 | AMGKEDYYDSSGPTGDY | 17 | 0 |  | 1-8 | 1 | QQYYSYPVT | 9 | 0 |  | - |
| aC2 s2p3 | 1-24 | 6-13 | 1 | 5 | ATDLMGSSWYQHWFDP | 16 | 0 |  | 1-39 | 3 | QQSYS | 5 | 0 |  | - |
| aC3 -S1P1 | 4-34 | D2-8 | 3 | 4 | ARGDVVYLYRH | 11 | 0 |  | 1-27 | 1 | QKYNSAPRT | 9 | 0 |  | + |
| aC3 S2P2 | 5-51 | D3-9 | 2 | 4 | ARSHYDILTGYYTAPFSPIDY | 21 | 0 |  | 2-28 | 1 | MQALQTPGT | 9 | 0 |  | - |
| aC4 -S1P1 | 3-23 | D3-3 | 2 | 4 | AKDIDYDFWSGSHGY | 15 | 0 |  | 3-11 | 1 | QQRSNWPGT | 9 | 0 |  | - |
| aD7 s2p1 | 4-59 | 2-21 | 3 | 3 | ARGGVVTAIGFRAIRDDAFDI | 21 | 0 |  | 3-15 | 1 | QQYNNWPP | 8 | 0 |  | + |
| aD9 s2p1 | 3-30 | 3-22 | 2 | 2 | ASDSRGNGLGHLVTFDL | 17 | 1 CDR3 |  | 3-15 | 5 | QQYNNWPIT | 9 | 0 |  | - |
| rIgG | VH | D | RF | JH | CDR3(aa) | Length | Mutations |  | Vλ | Jλ | CDR3(aa) | Length | Mutations |  | Poly |
| aC1 s2p3 | 4-34 | 2-15 | 3 | 4 | ARLLKVVAATRFDY | 14 | 0 |  | 1-44 | 3 | AAWDDSLNGPV | 11 | 1 CDR2 |  | - |
| aD11 s2p3 | 3-7 | 2-15 | 3 | 4 | ARELAVAAEEFDY | 13 | 1 FR3 |  | 2-8 | 1 | SSYAGSNSLYV | 11 | 0 |  | - |

RF: reading frame	-: non-reactive	+: reactive	FR: framework	CDR: complementarity-determining region	m: missense mutation.

Sequence and reactivity tables for new emigrant B cells from HC-1

|  | HEAVY | | | | | | |  | LIGHT | | | | |  | REACTIVITY |
| --- | --- | --- | --- | --- | --- | --- | --- | --- | --- | --- | --- | --- | --- | --- | --- |
| rIgG | VH | D | RF | JH | CDR3(aa) | Length | Mutations |  | Vk | Jk | CDR3(aa) | Length | Mutations |  | Poly |
| aE10 | 3-23 | 3-22 | 2 | 2 | AKDPKNYYDSSGYKNWYFDL | 20 | 0 |  | 4-1 | 2 | QQYYSTPL | 8 | 0 |  | - |
| aE11 | 3-30 | 1-26 | 4 | 2 | ARDPRRTRPELDY | 13 | 0 |  | 2-30 | 5 | MQGTHWPPIT | 10 | 0 |  | - |
| aE8 | 3-30 | 2-2 | 5 | 2 | ARNRYCSSTSCSYLPNWFDP | 20 | 0 |  | 3-15 | 3 | QQYNNWPHT | 9 | 0 |  | - |
| aE9 | 3-21 | 3-22 | 4 | 2 | ARDVSGLYYYDSSGYSGGD | 19 | 0 |  | 1-5 | 4 | QQYNSYFLT | 9 | 1 m CDR3 |  | - |
| aF11 | 3-48 | 2-15 | 5 | 2 | ARDCSGGSCYLNWFDP | 16 | 0 |  | 1-17 | 4 | LQHNSYPRS | 9 | 0 |  | - |
| aF5 | 3-23 | 4-17 | 4 | 2 | AKESYNYGDPVGVDY | 15 | 0 |  | 1-17 | 2 | LQHNSYPHT | 9 | 1 m CDR2 |  | - |
| aF6 | 4-4 | 2-15 | 2 | 5 | ARDGGWFDP | 9 | 0 |  | 1-39 | 1 | QQSYSTPRT | 9 | 0 |  | - |
| aF8 | 4-59 | 6-13 | 5 | 1 | ARRGSSWSAQNRDWFDP | 17 | 0 |  | 1-16 | 2 | QQYNSYPYT | 9 | 0 |  | - |
| aG1 | 3-23 | 6-13 | 4 | 2 | AKVALRMAAAGPFDY | 15 | 0 |  | 1-8 | 4 | QQYYSYPPT | 9 | 0 |  | - |
| aG10 | 1-8 | 6-13 | 4 | 1 | ARGGRYSRTFIAG | 13 | 0 |  | 3-15 | 2 | QQYNNWPYT | 9 | 0 |  | + |
| aG8 | 4-61 | 5-18 | 5 | 3 | ARERVSYGSYR | 11 | 0 |  | 3-20 | 2 | QQYGSSPYT | 9 | 0 |  | - |
| aH2 | 3-23 | 3-16 | 4 | 3 | AKVFFVVITFGGAIDY | 16 | 0 |  | 1-9 | 1 | QQLNSYPRT | 9 | 0 |  | + |
| aH9 | 3-23 | 3-22 | 4 | 2 | AETNYDSKLGGFDY | 14 | 0 |  | 3-15 | 1 | QQYNNWPRT | 9 | 1 m FR3 |  | - |
| aH4 | 4-38 | 2-15 | 2 | 5 | ARDATCSGGSCYSNWFDP | 18 | 0 |  | 1-39 | 1 | QQSYSTPWT | 9 | 1 m FR3 |  | - |
| rIgG | VH | D | RF | JH | CDR3(aa) | Length | Mutations |  | Vλ | Jλ | CDR3(aa) | Length | Mutations |  | Poly |
| aE2 | 3-23 | 3-22 | 6 | 2 | ASYYDSSGDYYYYGMDV | 17 | 0 |  | 3-27 | 2 | YSAADNKPELV | 11 | 1 m CDR1 |  | - |
| aF12 | 1-2 | 3-16 | 4 | 3 | ADMIRDY | 7 | 0 |  | 2-23 | 3 | CSYAGSSRV | 9 | 0 |  | - |
| aF9 | 3-7 | 4-17 | 4 | 2 | AREDNDYGDYFHY | 13 | 0 |  | 4-69 | 3 | QTWGTGIRV | 9 | 0 |  | - |
| aG4 | 3-23 | 6-19 | 4 | 1 | ASPLGVRGSRGWSACLGY | 18 | 0 |  | 3-21 | 3 | QVWDSSSDHRV | 11 | 0 |  | + |
| aG5 | 1-18 | 2-2 | 4 | 3 | ARDVFDIVVVPAYVRINPETLDY | 23 | 0 |  | 1-47 | 1 | AAWDDSLSGYYV | 12 | 0 |  | - |
| aG6 | 3-30 | 2-15 | 4 | 1 | ARDLYKWELPGRYRGFDY | 18 | 0 |  | 2-23 | 2 | CSYAGSSSVV | 10 | 0 |  | - |

RF: reading frame	-: non-reactive	+: reactive	FR: framework	CDR: complementarity-determining region	m: missense mutation

Sequence and reactivity tables for mature naïve B cells from HC-1

|  | HEAVY | | | | | | |  | LIGHT | | | | |  | REACTIVITY |
| --- | --- | --- | --- | --- | --- | --- | --- | --- | --- | --- | --- | --- | --- | --- | --- |
| rIgG | VH | D | RF | JH | CDR3(aa) | Length | Mutations |  | Vk | Jk | CDR3(aa) | Length | Mutations |  | Poly |
| aA8 | 4-39 | 1-26 | 4 | 1 | ARQVGATTIDY | 11 | 0 |  | 1-39 | 4 | QQSYSTPLT | 9 | 0 |  | - |
| aB11 | 1-2 | 3-9 | 3 | 3 | AREIFYVTSLWLPVGGFDI | 19 | 0 |  | 3-11 | 3 | QQRSNWPRVT | 10 | 1 m FR2, 1 m FR3 |  | + |
| aB12 | 3-23 | 5-18 | 4 | 1 | AKDEPDTAMVTLDY | 14 | 0 |  | 1-9 | 4 | QQLNSYPLT | 9 | 0 |  | - |
| aB7 | 3-23 | 2-21 | 4 | 3 | AKSVVAIDYFDY | 12 | 0 |  | 1-8 | 2 | QQYYSYPYT | 9 | 0 |  | - |
| aB9 | 3-23 | 5-24 | 4 | 2 | AKQRDEDYFDY | 11 | 0 |  | 1-5 | 1 | QQYNSYPGT | 9 | 0 |  | - |
| aC8 | 3-73 | 1-26 | 4 | 1 | TRQTIVGATTGDY | 13 | 0 |  | 1-12 | 4 | QQANSFPPT | 9 | 0 |  | - |
| aD11 | 3-73 | 4-17 | 4 | 3 | TRTTSFDY | 8 | 0 |  | 1-33 | 2 | QQYDNLPYT | 9 | 0 |  | - |
| aD12 | 1-2 | 3-22 | 4 | 2 | ARDLGPRDYDSSEGGFDY | 18 | 0 |  | 1-5 | 1 | QQYNSWWT | 8 | 0 |  | + |
| aD7 | 3-9 | 2-2 | 5 | 2 | AKAPGRTTRDWFDP | 14 | 0 |  | 3-15 | 4 | QQYNNWPPLT | 10 | 0 |  | - |
| rIgG | VH | D | RF | JH | CDR3(aa) | Length | Mutations |  | Vλ | Jλ | CDR3(aa) | Length | Mutations |  | Poly |
| aA2 | 3-48 | 2-2 | 4 | 3 | ARHSVPAAITLFDY | 14 | 0 |  | 3-10 | 2 | YSTDSSGNHRGV | 12 | 0 |  | - |
| aA6 | 3-21 | 1-26 | 6 | 1 | ARLGGAGEAPYYYYGMDV | 18 | 0 |  | 3-1 | 3 | QAWDSSNWV | 9 | 1 m FR2 |  | - |
| aB1 | 5-51 | 2-15 | 4 | 2 | ARGPPYCSGGSCYVFDS | 17 | 1 m CDR1, 1 m FR2, 1 m CDR2 |  | 2-14 | 1 | SSYTSSSTPYV | 11 | 1 m CDR1 |  | - |
| aC3 | 3-30 | 6-13 | 4 | 1 | AIFQNGGSSWTLGDY | 15 | 0 |  | 3-1 | 2 | QAWDSSTVV | 9 | 0 |  | - |
| aC7 | 3-9 | 6-13 | 4 | 3 | AKDLSGEQQLPGGY | 14 | 0 |  | 3-1 | 1 | QAWDSSTAI | 9 | 0 |  | - |
| aD2 | 1-69 | 3-22 | 5 | 3 | ALVVITTNWFDP | 12 | 0 |  | 8-61 | 3 | VLYMGSGISV | 10 | 0 |  | - |
| aD4 | 1-18 | 6-13 | 5 | 1 | ARESDAGYSSSWYDY | 15 | 0 |  | 3-1 | 2 | QAWDGSTVV | 9 | 1 m FR2, 1 m CDR3 |  | - |
| aD5 | 5-51 | 6-13 | 4 | 1 | ARRDGLGYSSSWCFDY | 16 | 0 |  | 2-23 | 2 | CSYAGSSTFGV | 11 | 0 |  | - |

RF: reading frame	-: non-reactive	+: reactive	FR: framework	CDR: complementarity-determining region	m: missense mutation
