## Supplementary figures and images for "B Cell Tolerance and BCR Signaling Dysregulation in NF155-Mediated Autoimmune Nodopathies"

### Supplementary Figures

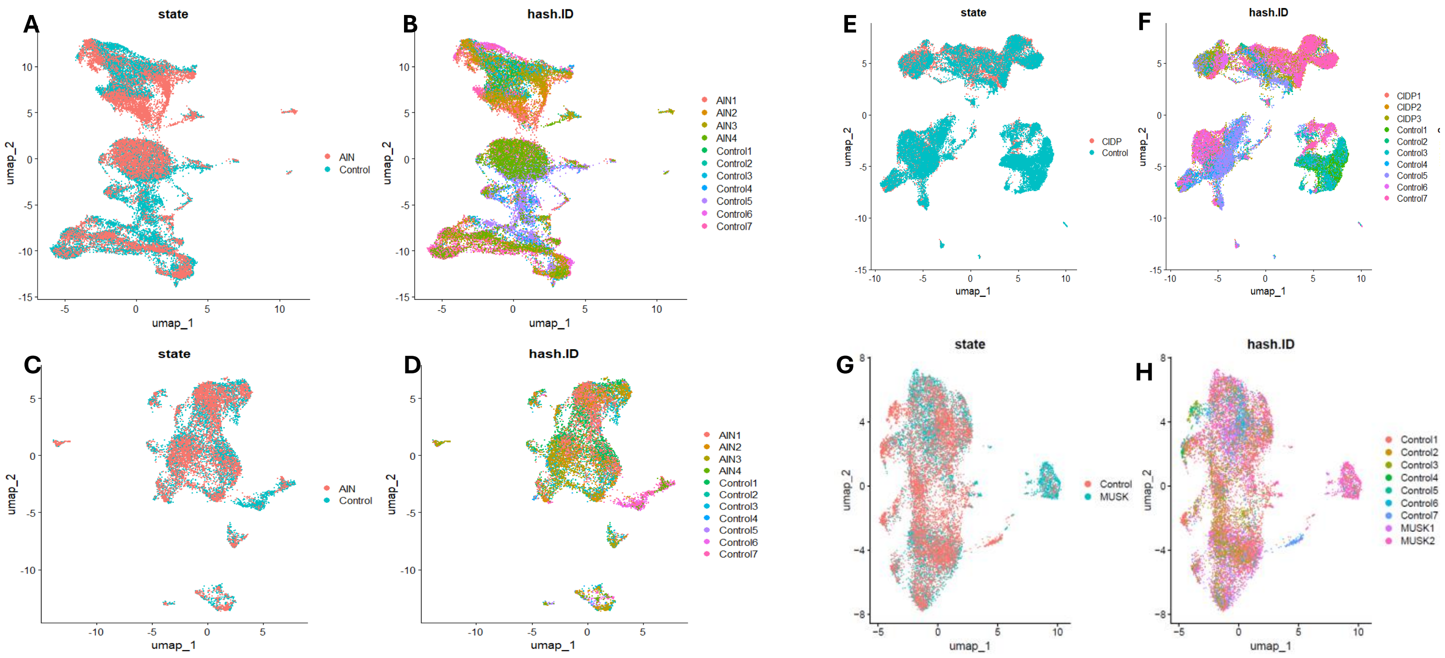

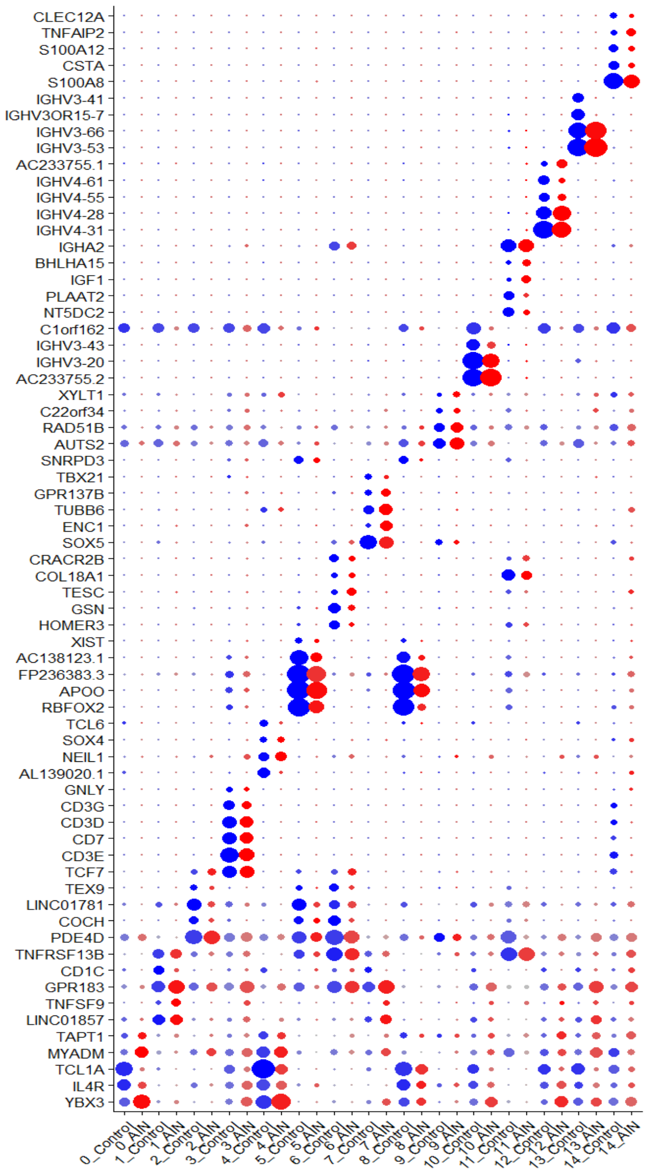

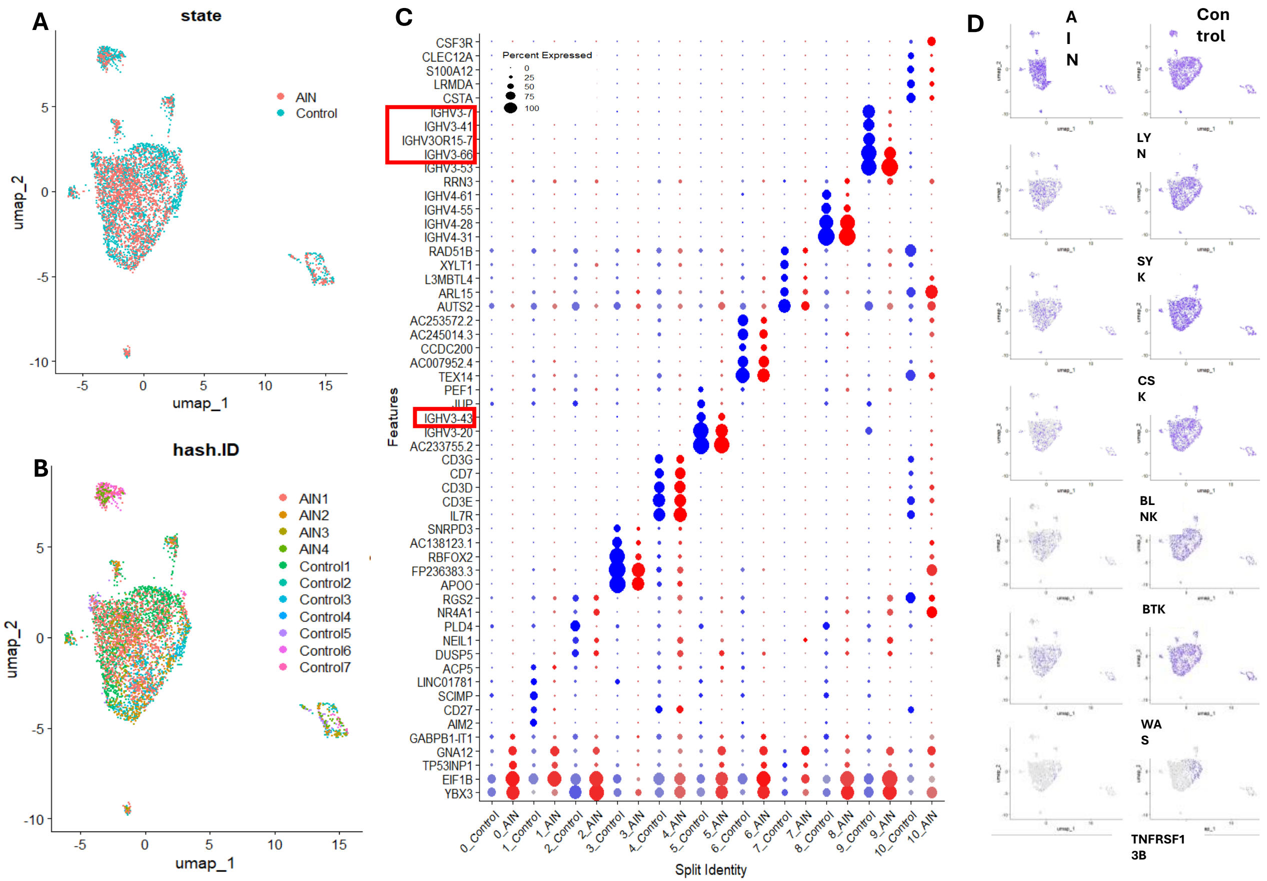

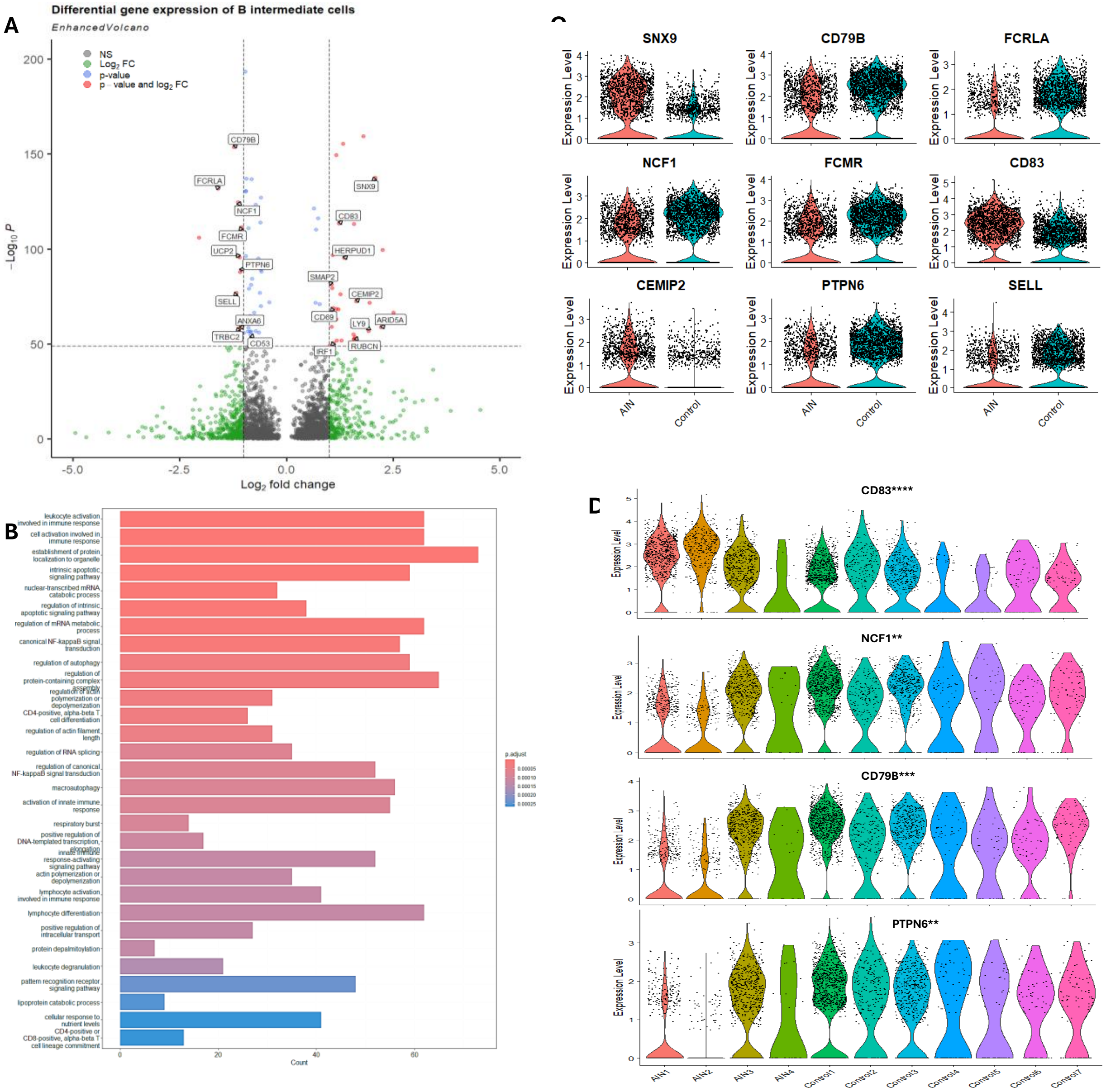

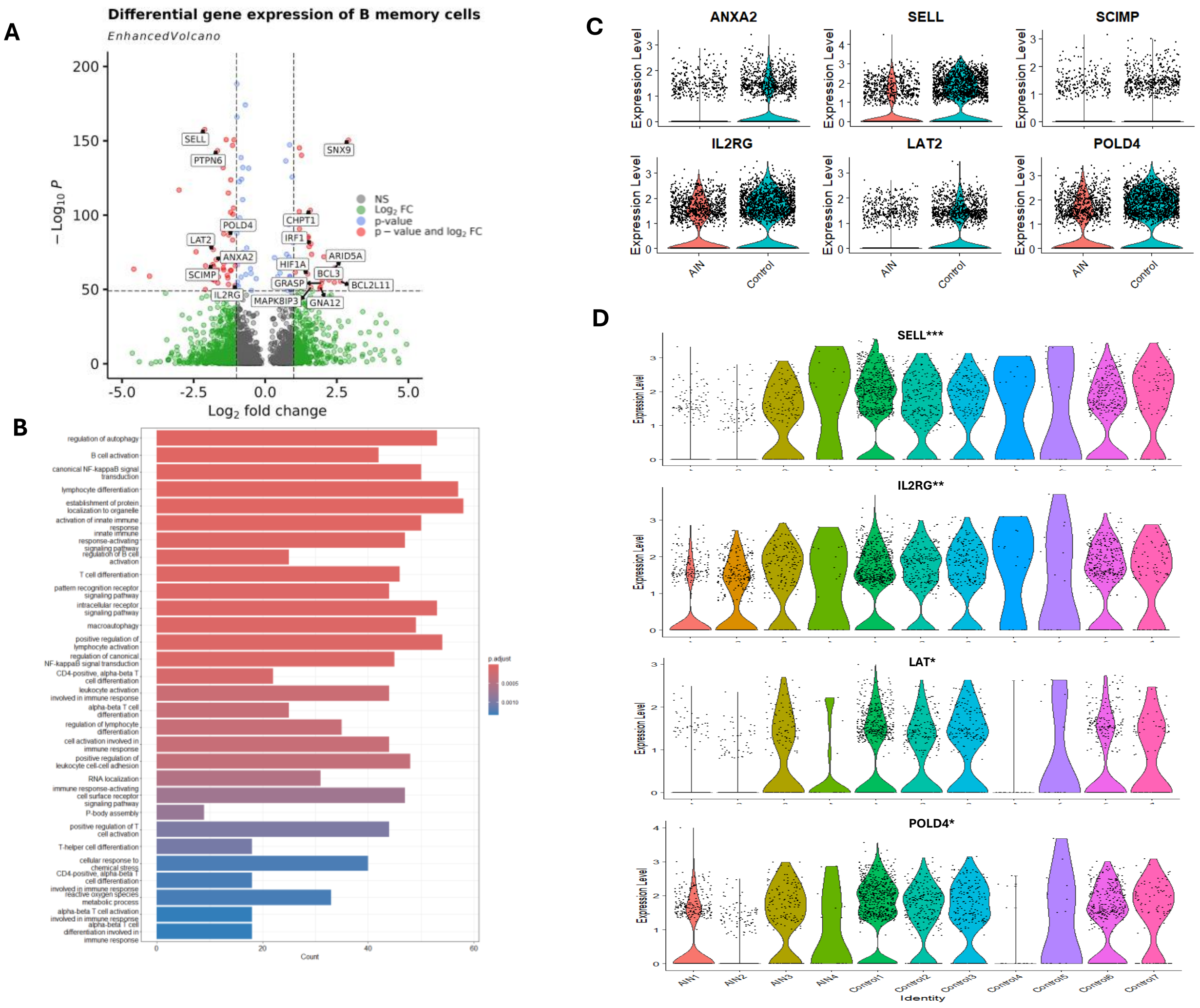

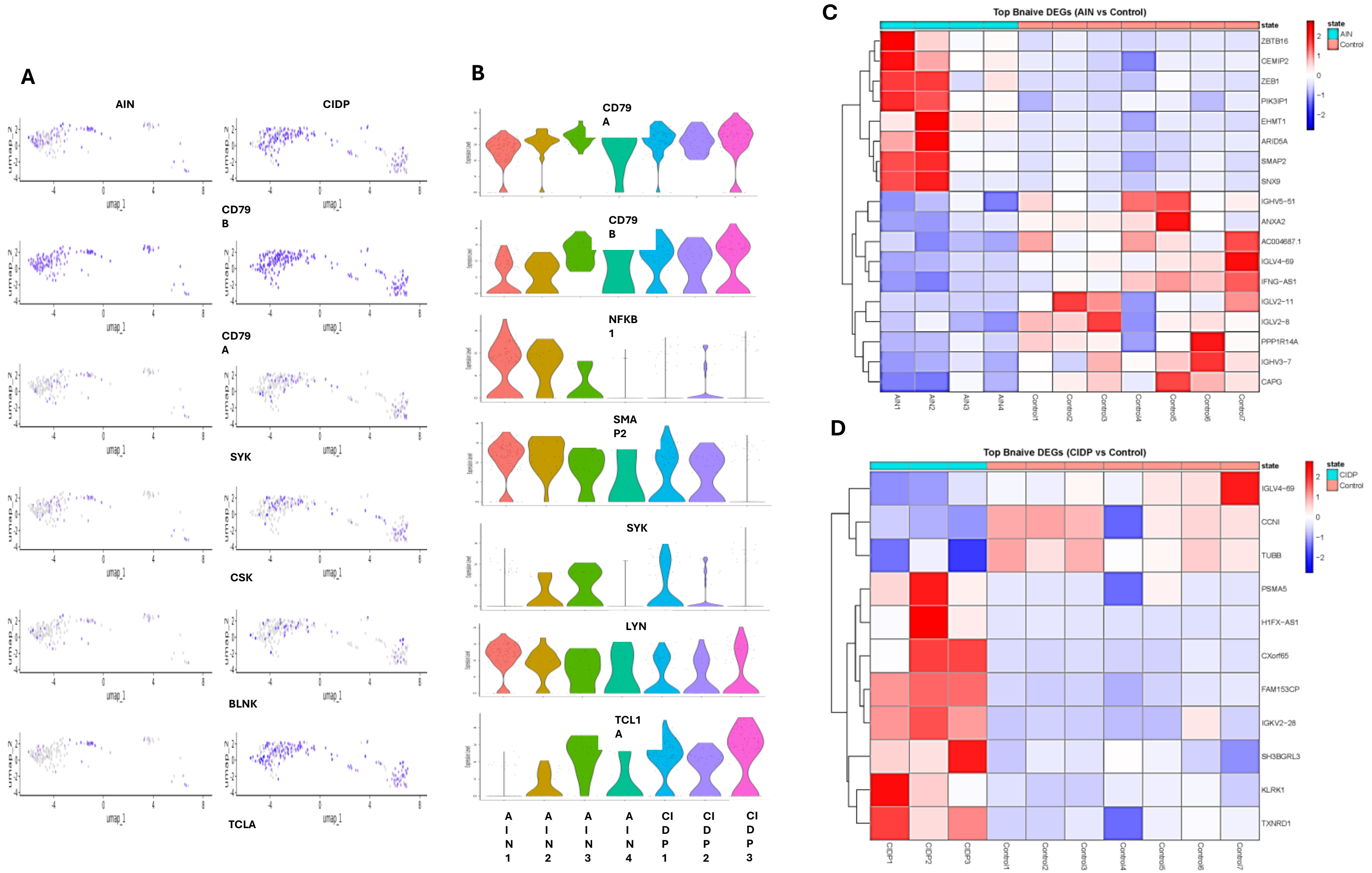

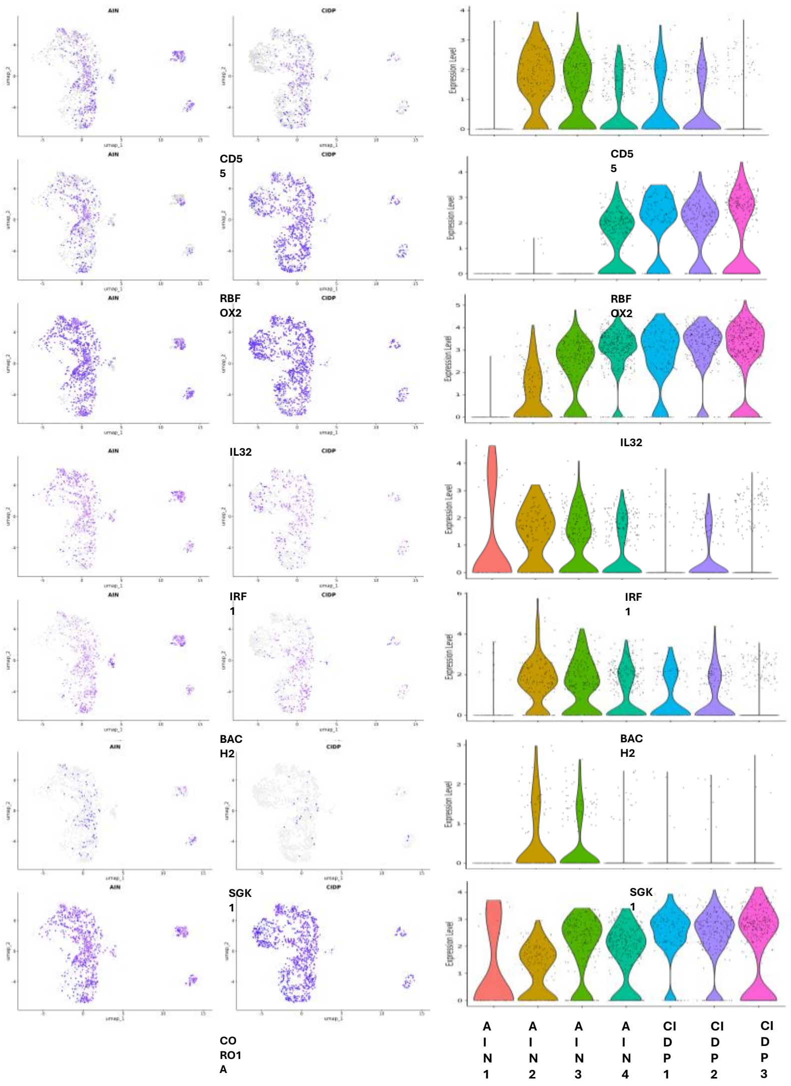

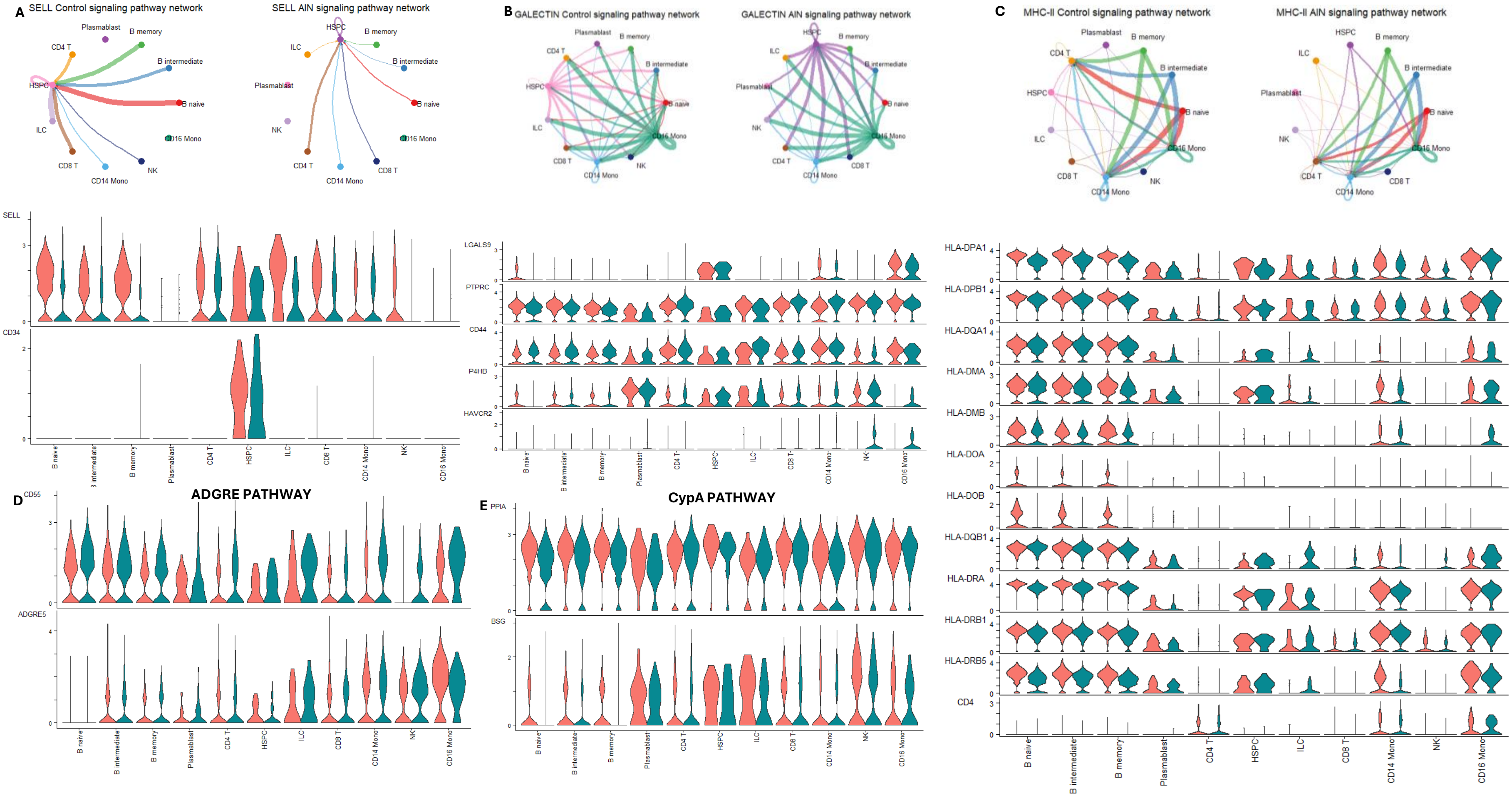
